# RNA polymerases reshape chromatin and coordinate transcription on individual fibers

**DOI:** 10.1101/2023.12.22.573133

**Authors:** Thomas W. Tullius, R. Stefan Isaac, Jane Ranchalis, Danilo Dubocanin, L. Stirling Churchman, Andrew B. Stergachis

**Affiliations:** Department of Genetics, Blavatnik Institute, Harvard Medical School, Boston, MA 02115, USA; Division of Medical Genetics, Department of Medicine, University of Washington, Seattle, WA; Department of Genome Sciences, University of Washington, Seattle, WA

## Abstract

During eukaryotic transcription, RNA polymerases must initiate and pause within a crowded, complex environment, surrounded by nucleosomes and other transcriptional activity. This environment creates a spatial arrangement along individual chromatin fibers ripe for both competition and coordination, yet these relationships remain largely unknown owing to the inherent limitations of traditional structural and sequencing methodologies. To address these limitations, we employed long-read chromatin fiber sequencing (Fiber-seq) to visualize RNA polymerases within their native chromatin context at single-molecule and near single-nucleotide resolution along up to 30 kb fibers. We demonstrate that Fiber-seq enables the identification of single-molecule RNA Polymerase (Pol) II and III transcription associated footprints, which, in aggregate, mirror bulk short-read sequencing-based measurements of transcription. We show that Pol II pausing destabilizes downstream nucleosomes, with frequently paused genes maintaining a short-term memory of these destabilized nucleosomes. Furthermore, we demonstrate pervasive direct coordination and anti-coordination between nearby Pol II genes, Pol III genes, transcribed enhancers, and insulator elements. This coordination is largely limited to spatially organized elements within 5 kb of each other, implicating short-range chromatin environments as a predominant determinant of coordinated polymerase initiation. Overall, transcription initiation reshapes surrounding nucleosome architecture and coordinates nearby transcriptional machinery along individual chromatin fibers.

## Introduction

Eukaryotic RNA Polymerase II (Pol II) transcription initiates in a highly regulated manner along a genome that is bound by numerous DNA-associated proteins including nucleosomes, transcription factors, chromatin remodelers, and other RNA polymerase complexes. Initiation begins with the assembly of the preinitiation complex (PIC) consisting of Pol II and an array of general transcription factors which work in unison to initiate transcription. Following PIC assembly and Pol II release, Pol II generally transcribes 25-50 bp and pauses at the promoter-proximal pause (PPP) site, a critical step in transcription that allows the integration of regulatory signals before proceeding into productive elongation^1,2^.

This chromatin structure creates a spatial arrangement conducive to both competition and coordination along individual chromatin fibers. Nucleosomes in particular serve as a physical barrier to both transcription initiation, pausing, and elongation^3–8^. Genes with higher rates of Pol II pausing are associated with distinct promoter-proximal nucleosome patterns^8^. However, it is unclear whether these patterns reflect paused Pol II directly modulating the positioning of assembled nucleosomes that it is competing with along an individual fiber. Further, transcription initiation occurs in spatial proximity to nearby transcriptionally active promoters and enhancers - a configuration that may promote coordination between these neighboring elements^9–11^. For example, neighboring promoters have correlated expression patterns, transcriptional activity at enhancers is associated with the expression of neighboring genes, and some transcriptionally active genes co-localize at specific foci within the nucleus^12–16^. It is unknown, though, if these associations represent direct, simultaneous coordination in transcription initiation along individual chromatin fibers within the nucleus.

Although pioneering cryo-EM and biophysical studies have provided a detailed single-molecule understanding of how transcription initiation occurs along individual DNA templates^4–6,17–20^, our understanding of this process along intact multi-kilobase chromatin fibers within the nucleus is limited. Emerging foot-printing methods^21–25^ have the potential to overcome these limitations, with single-molecule chromatin fiber sequencing (Fiber-seq) specifically providing near single-nucleotide resolution maps of protein occupancy along individual multi-kilobase fibers within the nucleus^22^. Fiber-seq uses a nonspecific *N*^6^-adenine methyltransferase (m6A-MTase) to stencil protein footprints onto their underlying DNA templates, which are directly read using PacBio SMRT sequencing, producing single-molecule maps of hundreds of individual protein footprints along 15-20 kb chromatin fibers^22^.

In this study we leverage Fiber-seq to directly characterize both competition and coordination between RNA polymerases and surrounding chromatin proteins along individual chromatin fibers. We present a hidden Markov model (HMM) based footprint caller (FiberHMM) that enables the identification and quantification of paused Pol II, elongating Pol II, PIC, nucleosome, and Pol III transcription-associated footprints. Using this single-molecule data we characterize the interplay between transcription initiation and its surrounding chromatin, including the mechanism for the inhibition of initiation by Pol II pausing. Further, we discover pause-driven changes to nucleosome architecture and direct coordination between nearby Pol II and Pol III genes, and transcribed enhancers within spatially organized chromatin domains.

## Results

### Identification of single-molecule RNA polymerase II footprints via FiberHMM

To identify single-molecule RNA polymerase-related footprints, we applied Fiber-seq to *Drosophila* S2 cells, resulting in an average sequencing coverage of 160x across the genome (12 kb average read length). Next, we developed a hidden Markov model (HMM) based footprint caller (FiberHMM) to identify unmethylated patches of adenines along each fiber that correspond to the footprints of RNA polymerase II, nucleosomes, and other chromatin-bound proteins. Specifically, FiberHMM incorporates observed adenine methylation patterns as well as false negative and false positive methylation probabilities for each base. These probabilities are derived from Fiber-seq data from methyltransferase-treated dechromatinized genomic DNA (gDNA) and methyltransferase-untreated genomic DNA, respectively. These false negative and positive methylation rates are used as fixed emission probability parameters within the HMM, while the starting and transition probabilities are trained using a subset of the Fiber-seq data. This trained model is then used to call gaps in methylation that represent protein footprints along each read at near single-base resolution **(Fig. 1a)**.

**Figure 1:**
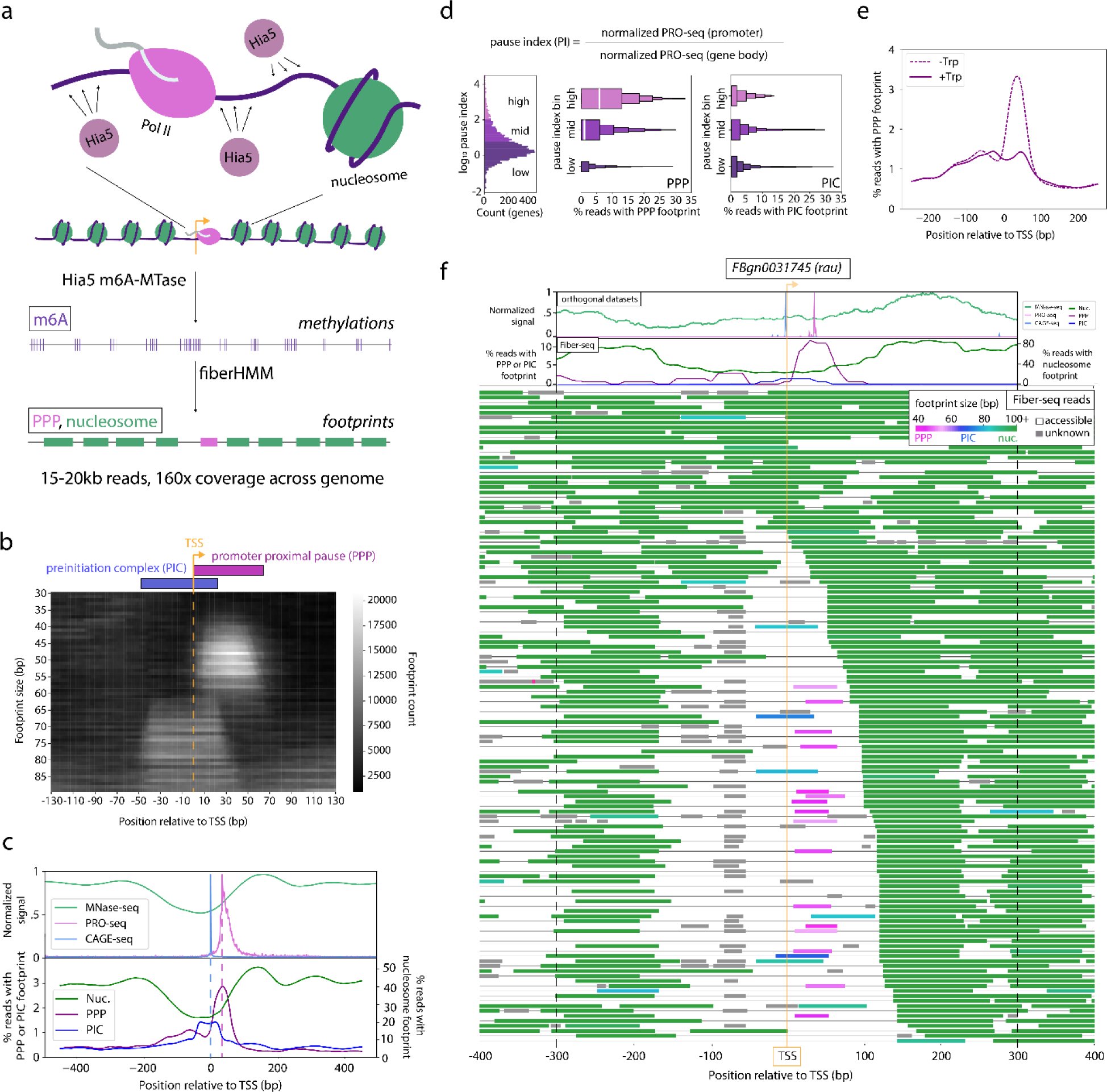
Identification of Pol II footprints in Fiber-seq. **(a)** Schematic of Fiber-seq. **(From top to bottom)** *D. melanogaster* S2 cells were subjected to nuclear permeabilization and treatment with Hia5 m6A methyltransferase. The resulting HMW-DNA were sequenced using PacBio SMRT sequencing, allowing direct detection of methylated bases. Footprints were then called using FiberHMM, resulting in ∼15-20 kb footprinted Fiber-seq reads representing individual chromatin fibers. **(b)** Heatmap depicting the enrichment of differently sized Fiber-seq footprints at positions around transcription start sites (TSS) of genes with a pause index ≥10. **(c)** Plot depicting the summed enrichment of **(top)** MNase-seq, PRO-seq, and CAGE-seq and **(bottom)** Nucleosome-, PPP-, and PIC-sized footprints at positions around TSSs of genes with a pause index ≥10. A dashed line is drawn from the peak of the CAGE-seq and PRO-seq peaks to visualize their alignment with the PPP and PIC footprints below. **(d)** A set of plots, showing a **(left)** histogram of the distribution of pause index values for genes binned by pause index: “high” (PI ≥ 100), “mid” (100>PI≥10), “low” (10>PI). Boxenplots showing the enrichment of (center) PPP and (right) PIC footprints in Fiber-seq reads from genes in each pause index bin. **(e)** Plot showing the enrichment of PPP-sized footprints in Fiber-seq in S2 cells (dashed) and S2 cells treated for 30 minutes with 10 µM triptolide (solid). **(f)** Fiber-seq reads mapped to an example locus. The top two tracks plot **(above)** MNase-seq, PRO-seq, and CAGE-seq signal, and **(below)** enrichment of nucleosome-, PPP-, or PIC-sized footprints. Individual Fiber-seq reads **(below)** are plotted on each line with footprints represented by colored blocks and accessible regions by a thin black line. Footprints are colored based on their size, with nucleosome-, PPP-, and PIC-sized footprints being green, pink, and blue respectively. Footprints smaller than a nucleosome, but not overlapping a PRO-seq or CAGE-seq peak, are colored gray to indicate that their identity is unknown. Reads are sorted by the position of the +1 nucleosome.

Promoter-proximal paused RNA polymerase II (PPP) and preinitiation complex (PIC) footprints have been previously observed using short-read MTase-based approaches^21^. As such, we sought to determine whether we could similarly identify PICs and PPPs among the FiberHMM-derived footprints at genomic sites known to harbor these complexes. When we aligned Fiber-seq footprints on a metagenomic scale around transcription start sites (TSSs), two distinct populations of footprints were visible: the first directly overlapping the TSS and a second located 0-60 bp downstream of the TSS **(Fig. 1b)**. This first footprint population matches the previously observed location of PICs, whereas the latter matches the previously observed location of PPPs. The size ranges of these putative PIC and PPP footprints ranged from 60 to 80 bp and from 40 to 60 bp respectively. Importantly, these ranges mirror the expected sizes from previous biochemical and structural studies^26–28^ **(Table S1)**.

### Validated single-molecule transcription complex occupancy

We next sought to validate the positioning and enrichment of these putative PIC and PPP footprints using a combination of orthologous datasets and chemical perturbations. First, we compared the positioning of these footprints to short-read sequencing derived transcript initiation and pausing data from CAGE-seq and PRO-seq, respectively^29–32^. As expected, putative PIC footprints are selectively positioned directly over CAGE-seq peaks, and putative PPP footprints are selectively positioned directly over PRO-seq peaks **(Fig. 1c)**. Second, we evaluated the enrichment of PPP footprints within TSSs for highly paused genes. As expected, putative PPP footprints were significantly enriched at genes with a high PRO-seq derived pause index, and gene pause index and PPP footprint enrichment showed a monotonic relationship **(Fig. 1d, Fig. S1c)**. In contrast, the enrichment of putative PIC footprints was independent of pause index **(Fig. 1d)** and the enrichment of putative PPP footprints was independent of expression **(Fig. S1a,b)**. Finally, we performed Fiber-seq on cells treated with triptolide, an initiation inhibiting compound that blocks progression from the PIC to the promoter proximal pause^33^. Treatment with triptolide significantly reduced the levels of putative PPP footprints, while having no effect on the levels of putative PIC footprints **(Fig. 1e, S1d,e)**.

In addition to PIC and PPP footprints, FiberHMM also identified nucleosome-sized footprints that are preferentially localized to sites that mirror bulk MNase-seq data, as well as smaller footprints upstream of the TSS that likely correspond to transcription factor (TF) occupancy events **(Figure 1f)**. Overall, each Fiber-seq read on average contains ∼70 protein occupancy events defined by FiberHMM, with the majority of these being nucleosome footprints, enabling us to evaluate the single-molecule co-dependency between PIC, PPP, and nucleosome occupancy at individual loci genome-wide **(Fig. 1f, Fig. S2)**.

### Pausing is associated with changes in nucleosome architecture

Genes with high pausing are known to be associated with disrupted chromatin patterns. Specifically, studies using MNase-seq have reported that highly paused genes tend to have decreased promoter accessibility and shifted upstream and downstream nucleosomes^7,8^. However, directly disentangling the relative contribution of promoter accessibility and PPP occupancy to these chromatin changes has been stymied by the lack of single-molecule resolution. Taking advantage of the ability of Fiber-seq to capture the occupancy of hundreds of proteins along a single chromatin fiber, we set out to determine the degree to which pause index-associated changes to nucleosome architecture are directly explained by paused Pol II itself.

In aggregate, we found that Fiber-seq footprints mirrored the MNase-seq derived patterns of nucleosome enrichment seen at genes with different pausing and expression levels **(Fig. S3a)**^34^. Similarly, we found that genes with higher levels of pausing had more fibers with inaccessible promoters, in contrast to more highly expressed genes, which had more fibers with accessible promoters **(Fig. S3b)**. Importantly, we observed that chromatin fibers with inaccessible versus accessible promoters had markedly distinct nucleosome architectures, which are unavoidably aggregated together when analyzing MNase-seq data **(Fig. S3c)**. To remove the impact inaccessible promoters have on the overall pattern of nucleosome enrichment, we selectively evaluated nucleosome patterns on chromatin fibers containing accessible promoters. This analysis demonstrated that genes with high pause indices are, in fact, associated with shifted +1 and −1 nucleosomes **(Fig. 2a)**. In contrast, controlling for promoter accessibility, we found no clear difference in nucleosome architecture based on expression levels of genes, suggesting a unique role for pausing in altering the nucleosome landscape **(Fig. S3d)**.

**Figure 2:**
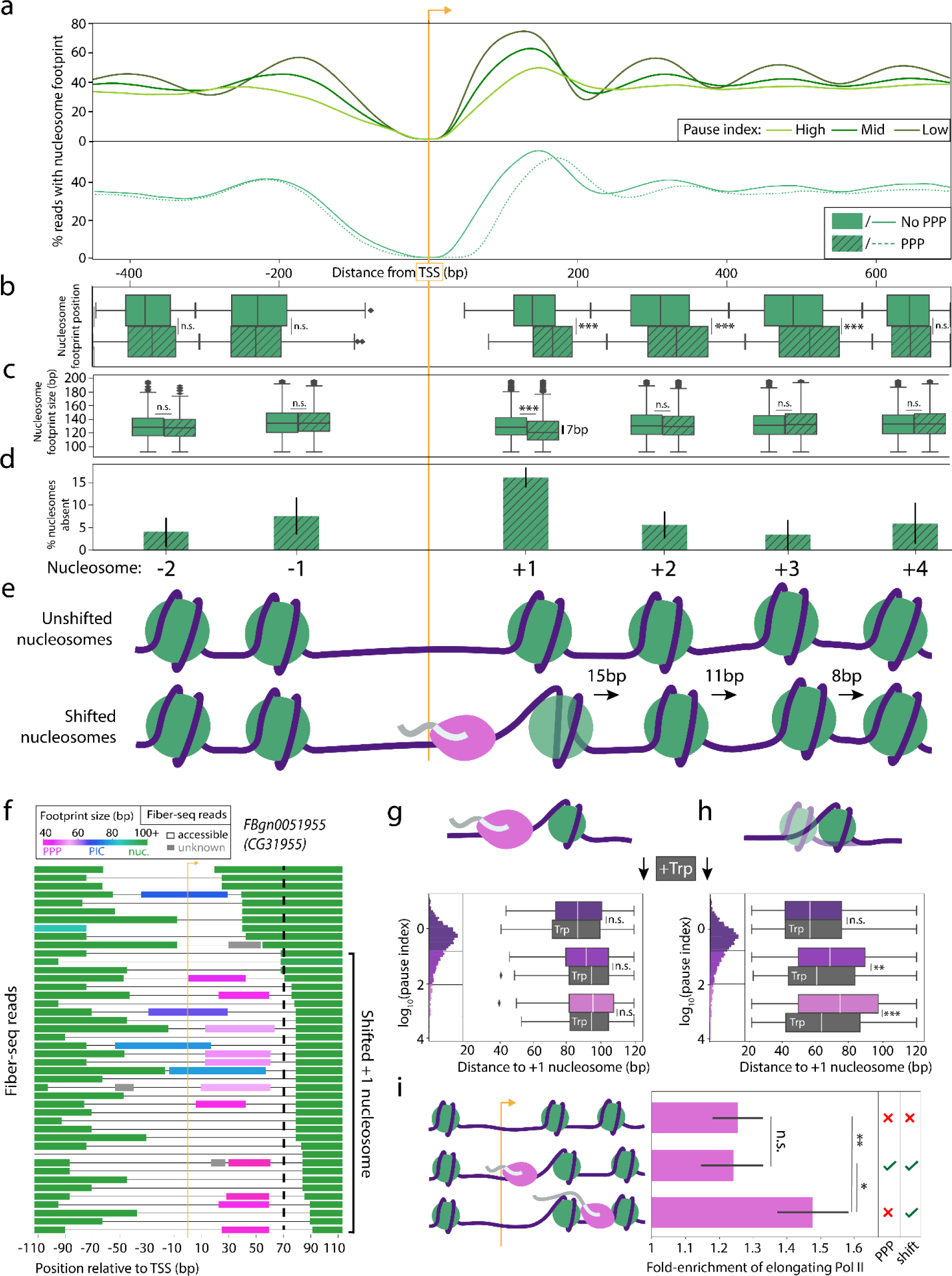
Pausing drives changes to downstream nucleosomes. **(a) (Top)** Enrichment of nucleosome footprints from genes binned by pause index: “high” (PI ≥ 100), “mid” (100>PI≥10), “low” (10>PI). Reads are filtered to only include those without a nucleosome overlapping the TSS. **(Bottom)** Plot depicting the enrichment of nucleosome footprints from reads with (solid) or without (dashed) a PPP footprint. Reads are filtered to those with an accessible promoter and are sampled to include an equal count of reads with or without a PPP footprint from each gene. **(b)** Boxplot showing the distribution of the position of each nucleosome upstream and downstream of the TSS between reads with (dashed) or without (solid) a PPP footprint. The nucleosome footprints were segmented using a Gaussian mixture model (GMM) based on a 95% posterior probability, and the distributions were compared with a two-sample t-test (n.s. signifies p>.05, *** signifies p-value <10^−13^). **(c)** Boxplot showing the size distribution of nucleosomes from reads with (dashed) or without (solid) a PPP footprint. Size distributions were compared using a two-sample t-test (n.s. signifies p>.05, *** signifies p-value < 10^−14^). **(d)** Bar plot showing the percentage fold-increase of absent nucleosome footprints in reads with a PPP footprint compared to reads without a PPP footprint. Error bars represent the confidence interval derived from the bootstrapped distribution of the ratio of reads with or without a nucleosome footprint at each nucleosome position in reads with or without a PPP footprint. **(e)** Cartoon summarizing the changes to nucleosome position and stability associated with the PPP. **(f)** Sample of Fiber-seq reads from a representative locus (CG31955), with footprints colored based on predicted identity (PPP = pink, PIC = blue, nucleosome = green, unknown = gray). A line is added to indicate reads with the shift of the +1 nucleosome described above, defined as starting between 70 and 150 bp downstream of the TSS. **(g, h)** Boxplots showing the distance to the +1 nucleosome footprint across reads with **(g)** or without **(h)** a PPP footprint. Reads are binned by pause index, as schematized in the associated histogram: “high” (pink, PI ≥ 100), “mid” (purple, 100>PI≥10), “low” (dark purple, 10>PI). +1 nucleosome distances from cells treated with triptolide are plotted in gray boxplots below the corresponding colored box plots corresponding to −Trp cells. Significance is indicated between +Trp and −Trp within each pause index bin. (two-sample t-test, n.s. signifies p>.05, * signifies p <.05, ** signifies p < .01, *** signifies p < .0001). **(h)** Bar plot showing the fold enrichment of putative elongating Pol II footprints within gene bodies compared to the level of equivalently sized footprints in intergenic regions, divided into reads with no PPP and no +1 nucleosome shift **(top)**, a PPP and a +1 nucleosome shift **(middle)**, or no PPP and a +1 nucleosome shift **(bottom)**. Significance of enrichment of elongating Pol II footprints is indicated between each pair of sets of reads (two-sample t-test, n.s. signifies p>.7, * signifies p-value < .03, ** signifies p-value < 0.007).

Based on this analysis alone, though, we could not rule out if these changes to nucleosome architecture simply reflected sequence differences of highly paused genes or the impact of the PPP itself. To specifically identify PPP-associated changes, we further divided Fiber-seq reads containing an accessible promoter based on whether they also contained a PPP footprint. We then compared the positioning of surrounding nucleosome footprints for each fiber type. The proportion of accessible fibers with a PPP footprint varies genome-wide and are in the minority at all promoters **(Fig. 1d)**. To ensure that each promoter contributes the same number of fibers with and without a PPP, we subsampled an equal number of each fiber type at individual promoters. We found that PPP footprints across all genes were associated with disrupted down-stream nucleosome positioning, similar to the global effects seen associated with pause index **(Fig. 2a)**. However, no significant difference was observed for upstream nucleosomes, indicating that the change in the −1 nucleosome position was independent of the PPP. This strong directionality suggested an active role for the PPP in defining the nucleosome landscape.

Overall, we observed that chromatin fibers containing a PPP footprint had multiple changes in the organization of the +1 to +4 nucleosomes. First, the +1 nucleosome was shifted downstream by ∼15 bp and had a footprint ∼7 bp smaller, indicating that this nucleosome is displaced and often either partially unwound or a hexasome **(Fig. 2b-c**, **Fig. 4a-d)**. The +2 and +3 nucleosomes also were significantly shifted down-stream, ∼11 bp and ∼8 bp respectively, but showed no significant size difference **(Fig. 2b-c)**. Second, we observed that some chromatin fibers containing a PPP footprint were missing the +1 nucleosome, indicating that PPP footprints are associated with displacement of downstream nucleosomes **(Fig. 2d)**. In contrast, these effects were not seen upstream of the promoter or further than 1 kb downstream of the promoter and were similarly not seen in response to PIC occupancy within the promoter **(Fig. 2a-d, Fig. S3e, Fig. S4e-f)**. Taken together, these observations illustrate a nearby downstream nucleosome landscape transiently and significantly altered in association with promoter proximal pausing **(Fig. 2e)**.

### Pause-associated disruption of downstream nucleosomes is stably maintained

We next determined whether these altered downstream nucleosomes could be directly attributed to promoter proximal pausing. Notably, at highly paused genes, we found an increase in the frequency of fibers with a shifted +1 nucleosome, even on fibers without a PPP footprint, showing that the +1 nucleosome shift can exist independently of PPP co-occupancy **(Fig. 2f,h)**. However, at genes with a low pause index, the shifted +1 nucleosome was primarily seen only on reads with a PPP **(Fig. 2g)**. Together, these findings suggest a model whereby promoter-proximal pausing directly causes the +1 nucleosome to shift, which is then maintained for a brief but detectable period, and is thus visible only at loci with strong levels of pausing.

To test this model, we performed Fiber-seq after significantly reducing the formation of PPPs using triptolide. As described earlier, triptolide treatment markedly reduced the proportion of reads containing a PPP footprint, with the few reads that did contain a PPP footprint showing the expected downstream shift of the +1 nucleosome **(Fig. 2g)**. In contrast, the reads that lacked a PPP footprint had a dramatically reduced shift in the +1 nucleosome **(Fig. 2h)**. This reduction in the shift of the +1 nucleosome was particularly strong at highly paused genes **(Fig. 2h)**, which also showed the largest net reduction in PPP footprints. Notably, the +1 nucleosome distance at highly paused genes now mirrored the distances observed on lowly paused genes in the untreated sample. Thus, the distinct positioning of the +1 nucleosome at highly paused genes depends on the level of pausing. Together, these data demonstrate that promoter proximal pausing is directly linked to a shift in the +1 nucleosome and that this shifted +1 nucleosome is maintained on that chromatin fiber after the PPP has been released.

### Disrupted downstream nucleosomes are associated with active transcription

We hypothesized that if the reads with a shifted +1 nucleosome and no PPP footprint represented a transient state after pausing, they would be enriched for fibers with a post-pause released Pol II in the midst of productive gene-body elongation. To test this, we first established whether Fiber-seq could identify elongating RNA Pol II footprints, which we expected to be a similar size to the PPP given their similar chromatin contacts and structure^35,36^. Across all genes and reads, we found putative elongating Pol II footprints (i.e., 40-60 bp footprints within a gene body) at a frequency of ∼1.5 per 10 kb on average, which was markedly enriched at highly transcribed genes **(Fig. S5a)**. Furthermore, elongating Pol II footprints were depleted on gene bodies downstream of inaccessible promoters **(Fig. S5b)** and in cells treated with triptolide **(Fig. S5c)**, suggesting that the density of elongating Pol II footprints corresponds to the level of gene body transcription.

Next, we set out to determine if reads with a shifted +1 nucleosome, but no PPP footprint, represented fibers enriched for productive gene body transcription. Compared to reads without a recently released PPP (i.e., an unshifted +1 nucleosome, or a PPP footprint), reads with a recently released PPP (i.e., no PPP and a shifted +1 nucleosome) had a significantly higher density of elongating Pol II footprints **(Fig. 2i)**. Consequently, we conclude that Pol II pausing establishes a state of disrupted downstream nucleosomes, which upon pause release is maintained and is associated with transcriptional elongation.

### Pausing sterically inhibits initiation

Having demonstrated coordination between pausing and the downstream nucleosome landscape, we next sought to address how pausing and upstream initiation are coordinated within individual promoter regions. It is well established that pausing inhibits initiation, which is proposed to be explained, at least in part, through the steric interference between the PPP and PIC complexes^17,37,38^. To directly test this model, we inspected the frequency of PIC and PPP footprints within the same promoter region. Overall, we found that PPPs and PICs rarely co-occurred on the same chromatin fiber **(Fig. 3a)**. Notably, chromatin fibers containing simultaneous PPP and PIC footprints were observed as frequently as expected at loci with primary CAGE-seq and PRO-seq peaks separated by more than 60 bp, and especially more than 75 bp, a distance that would allow for unobstructed binding of both complexes **(Fig. 3b)**. In contrast, genes with primary CAGE-seq and PRO-seq peaks separated by fewer than 60 bp were significantly depleted in chromatin fibers with simultaneous PPP and PIC footprints **(Fig. 3b)**. This strong distance-dependent effect supports steric interference as the primary mechanism of pause-inhibited initiation.

**Figure 3:**
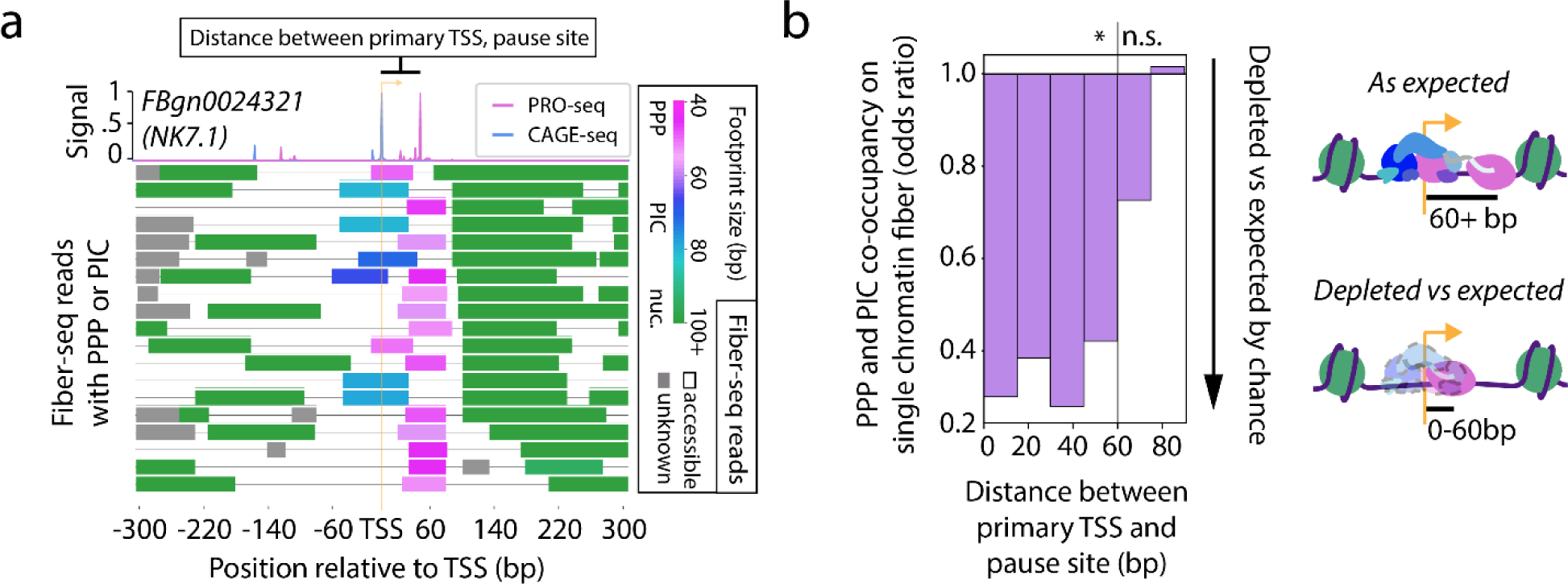
Pausing sterically inhibits initiation. **(a)** Sample Fiber-seq reads from a locus with simultaneous PPP and PIC footprints, with footprints colored based on predicted identity (PPP = pink, PIC = blue, nucleosome = green, unknown = gray). **(top)** Corresponding PRO-seq (pink) and CAGE-seq (blue) signal at the locus. **(b)** Bar plot, showing the Fisher’s exact test odds ratio of PPP and PIC footprint co-occupancy on a single read. Genes are binned based on the distance between their primary (tallest) PRO-seq and CAGE-seq peaks. Significance is indicated above the plot (Fisher’s exact test, n.s. signifies p > .05, * signifies p-values ranging from p < .05 to p < 10^−25^).

### Coordinated transcription initiation at neighboring genes

RNA expression studies have shown that transcription at nearby pairs of genes is often correlated in *Drosophila* and other eukaryotes, with this effect predominantly seen at gene pairs located less than 1 kb from each other^9,10,39^. Further, imaging studies have demonstrated that Pol II localization within the nucleus is often clustered^12–16^, suggesting a model whereby spatially proximal pairs of genes may obtain correlated expression via the coordinated occupancy of spatially clustered promoters. However, directly testing this model requires simultaneously observing transcription initiation events at single-molecule resolution across multi-kilobase chromatin fibers genome-wide, which was possible using Fiber-seq.

We next set out to assess whether transcription influences activity at the promoters of neighboring genes. First, we asked whether spatially proximal transcribed gene promoters were preferentially accessible along the same chromatin fiber and found that they are, suggesting coordination in the chromatin architectures at neighboring gene promoters **(Fig. S6a)**.

To directly test whether transcription initiation itself is coordinated between the nearby promoters of different genes, we evaluated the frequency of co-occupancy by transcription initiation footprints at all pairs of promoters captured within a single Fiber-seq read. We specifically included only reads from chromatin fibers on which both promoters were accessible, thereby accounting for the effect of promoter co-accessibility. Overall, we found that reads overlapping pairs of accessible promoters located within 7.5 kb of each other were preferentially co-occupied by PPP and PIC footprints significantly more frequently than expected. Furthermore, we observed a strong distance-dependent relationship to this effect, with the strongest co-occupancy seen at paired promoters located within 1 kb of each other **(Fig. 4a)**. These global effects were recapitulated on a per-gene basis **(Fig. S6c-d)**. Together, these findings indicate that correlated expression of spatially proximal gene pairs is driven in part by the coordinated occupancy of RNA polymerase complexes at both promoters simultaneously.

**Figure 4:**
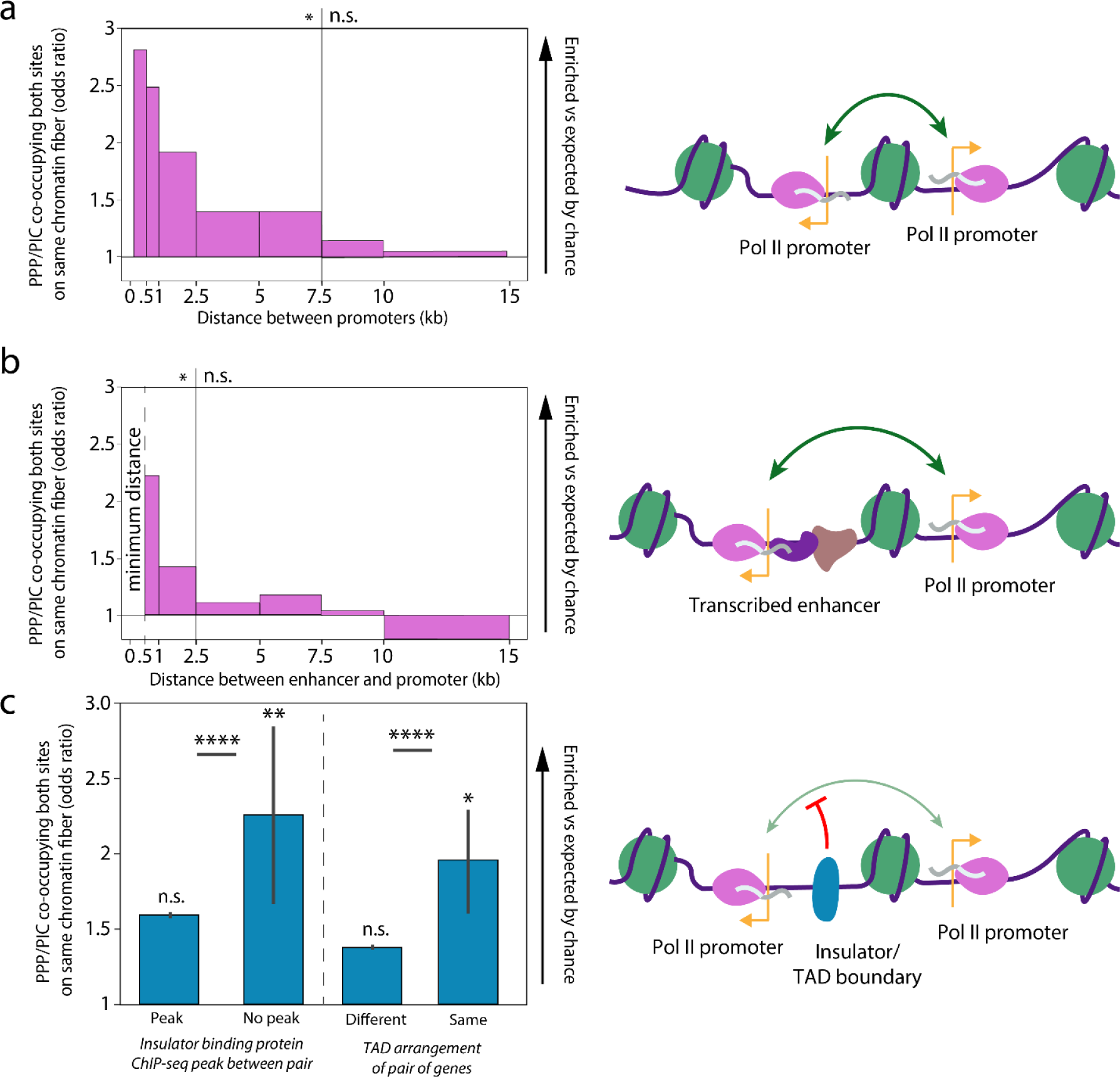
Transcription initiation is coordinated based on proximity. **(a)** Bar plot showing the Fisher’s exact test odds ratio of PPP/PIC footprint co-occupancy on reads over-lapping promoter pairs binned by distance. Significance is indicated above the plot. (Fisher’s exact test, n.s. signifies p > .05, * signifies p-values ranging from p < .001 to p < 10^−12^). **(b)** Bar plot distance, showing the Fisher’s exact test odds ratio of PPP/PIC footprint co-occupancy on reads overlapping enhancer-promoter pairs binned by distance. Significance is indicated above the plot. (Fisher’s exact test, n.s. signifies p>.05, * signifies p-values ranging from p < .001 to p < 10^−9^) at up to 5 kb separation. Minimum distance indicated on the plot indicates that enhancers were defined as at minimum 500bp from a known TSS. **(c)** Bar plot depicting the coordination of transcription initiation at **(left)** pairs of genes in different or the same topologically associating domain (TAD) and **(right)** pairs of genes separated or not separated by a ChIP-seq peak of an insulator binding protein. All pairs are found within 2.5 kb of each other and reads from each pair of groups are sampled to capture an identical count of reads from pairs at any given distance, with error bars corresponding to the confidence interval calculated from 10,000x sampling iterations. Pairwise significance is indicated between each set of comparisons (two-sample t-test, **** signifies p-value < 10^−193^), and mean significance for each individual sample across all sampling iterations is indicated (Mean of sample Fisher’s exact test results, n.s. signifies p > .05, * signifies p < .05).

### Coordinated transcription initiation between enhancers and nearby genes

Many eukaryotic enhancers are transcribed by RNA polymerase II to form enhancer RNAs (eRNAs), the production of which is strongly associated with enhanced transcription at nearby genes^40–42^. Given the coordination of transcriptional machinery between neighboring protein-coding gene promoters, we hypothesized that transcription initiation at transcribed enhancer-promoter pairs may be similarly coordinated. Using *Drosophila* S2 cell STARR-seq and START-seq data, which map enhancer regions and nascent transcription start sites, respectively^40,43,44^, we identified ∼4,000 enhancers that produce eRNAs. Overall, eRNA PPP and PIC footprints mirrored those at coding promoters, consistent with them being initiated by RNA polymerase II **(Fig. S7, Fig. S8a-e)**. Like paired coding promoters, we found that chromatin fibers overlapping eRNA-producing enhancers and a neighboring promoter were preferentially co-occupied by initiating polymerase footprints significantly more frequently than expected based on the overall frequency of the footprints. Even more so than in the case of coding promoters, we found a strong distance-dependent relationship in this effect **(Fig. 4b)**. Together, these findings suggest that the proximity of an eRNA-producing enhancer adjacent a promoter enables the coordinated occupancy of RNA polymerase complexes at both simultaneously, thereby potentiating the transcription of both the enhancer and gene.

### Coordinated initiation is enriched within TADs and blocked at TAD boundaries

Given the importance of 3D genome architecture in spatially connecting promoter and enhancer activity^11,43–45^, we set out to determine if the coordination in transcription initiation between pairs of genes based on 2D proximity would be affected by 3D chromatin architecture. Eukaryotic genomes are partitioned into topologically associating domains (TADs)^46–50^. We divided the pairs of genes from the previous coordination analysis based on if they were both contained in the same TAD, or if they were located in separate TADs based on Hi-C data^50^. Controlling for differences in the distribution of inter-promoter distances in the same-TAD and different-TAD groups, we found that fibers overlapping pairs of genes split by TAD boundaries had significantly less evidence of transcriptional coordination **(Fig. 4c)**. TAD boundaries are generally defined in *Drosophila* by insulator proteins including CP190, CTCF, SuHW, and Mod, and we observed that pairs of genes separated by the binding sites of these factors showed significantly lower than expected transcriptional coordination **(Fig. 4c, S6b)**^51–53^. Together, these results demonstrate that coordination is disrupted by TAD boundaries and insulator elements, demonstrating an additional layer of regulation beyond proximity.

### Identification of RNA polymerase III-associated footprints via FiberHMM

We next sought to determine whether this coordination was a feature of other RNA polymerases, specifically RNA polymerase III. RNA polymerase III is primarily responsible for the transcription of tRNA and 5S rRNA genes, which are often highly expressed and spatially clustered together along the genome^54^, and are traditionally challenging to uniquely study using short-read sequencing-based approaches owing to their high sequence similarity. We first sought to determine whether FiberHMM could uniquely identify footprints at tRNA genes associated with RNA polymerase III transcription, which is mediated by TFIIIB and TFIIIC occupancy at specific sequence elements upstream of and within the tRNA gene^55,56^. Consistent with our expectations, we identified several distinct populations of footprints enriched at tRNA genes corresponding to the known binding sites of TFIIIB and TFIIIC **(Fig. 5a)**. Additionally, the size and position of these putative Pol III transcription associated footprints was consistent across all tRNA genes **(Fig. 5b)**.

**Figure 5:**
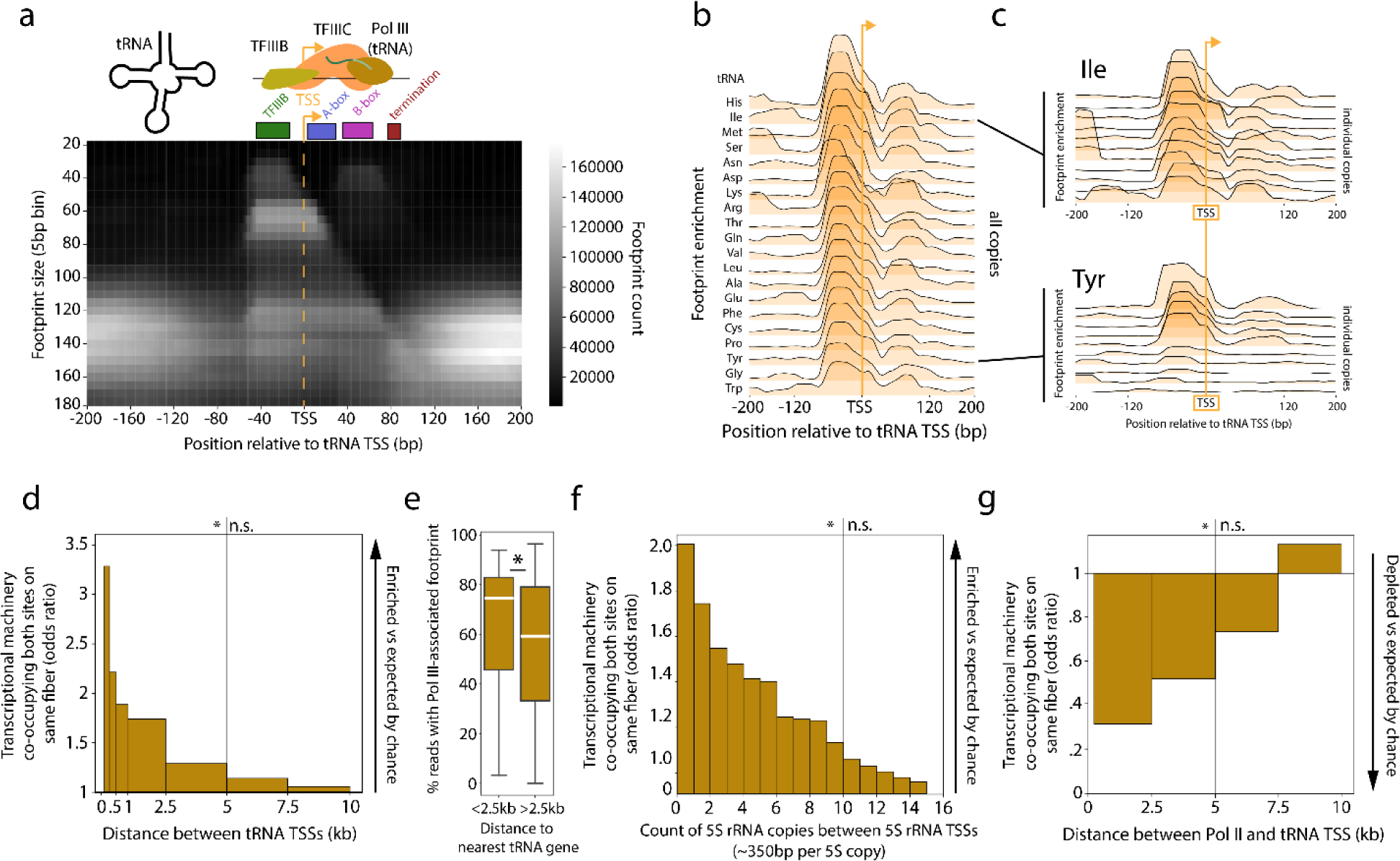
Coordination in transcription activity between nearby Pol III genes. **(a)** Heatmap depicting the enrichment of differently sized Fiber-seq footprints with respect to the nearest tRNA TSS. The expected position of TFIIIB, TFIIIC, and terminating Pol III are schematized as well as the A-box and B-box. **(b)** Set of plots showing the percent enrichment of 30-90 bp footprints in the region surrounding the TSS for each family of tRNA genes with a given amino acid, with families sorted by maximum percent enrichment. **(c)** Enrichment of 30-90 bp footprints relative to the TSS for individual isoleucine **(top)** or tyrosine **(bottom)** tRNA genes. Genes are sorted by maximum percent enrichment. **(d)** Bar plot depicting the Fisher’s exact test odds ratio of Pol III transcription footprint co-occupancy between pairs of tRNAs binned by distance. Significance is schematized above the plot (Fisher’s exact test, n.s. signifies p > .05, ** signifies p-value < 10^−4^, **** signifies p-value < 10^−85^). **(e)** Box plot showing the distribution of transcription frequency scores for tRNA genes either < 2.5 kb separated, or > 2.5 kb separated from another tRNA gene. Pairwise significance is indicated on the plot (two-sample t-test, * signifies p-value < 0.05). **(f)** Bar plot depicting the Fisher’s exact test odds ratio of Pol III 5S rRNA transcription associated footprint co-occupancy between pairs of 5S rRNAs binned by count of 5S rRNA copies separating the pair. Significance is schematized above the plot (Fisher’s exact test, n.s. signifies p > .05, ** signifies p-value < 10^−4^, **** signifies p-value < 10^−251^). **(g)** Bar plot depicting the Fisher’s exact test odds ratio of Pol III tRNA transcription associated footprint co-occupancy with PPP/PIC footprints between pairs of tRNA and Pol II genes binned by distance, horizontally scaled on distance. Significance is schematized above the plot (Fisher’s exact test, n.s. signifies p > .05, * signifies p-value < .05, ** signifies p-value < .01).

We next sought to determine whether the Pol III transcription associated footprints are related to tRNA expression. However, determining the expression level of individual tRNA genes using transcript-based sequencing is difficult as most tRNA genes in *Drosophila* have multiple copies that are identical in sequence. Furthermore, tRNA transcripts form strong secondary structures and undergo numerous modifications that impair standard reverse transcription-based transcript profiling methods^57–60^. Notably, we observed that individual copies of the same tRNA gene can have drastically different levels of these Pol III transcription-associated footprints, suggesting a high degree of variability in transcription activity between tRNA genes that produce sequence-identical transcripts **(Fig. 5c, Fig. S9a-c)**. To determine whether this variability in footprint occupancy reflected the transcriptional output of each tRNA gene, we calculated a ‘transcription frequency’ for each tRNA gene, which corresponds to the fraction of reads mapping to that gene occupied by Pol III transcription associated footprints. We found that the sum of these scores for all tRNA genes that contribute to the same codon significantly correlated with frequency of that codon within the *Drosophila* genome^61^. This correlation indicated that the transcriptional activity score of a tRNA gene accurately captured the demand for that tRNA within *Drosophila* cells **(Fig. S9d)**.

### Clustered Pol III genes exhibit coordinated transcription activity

As tRNA genes are often spatially clustered along the genome, we next tested whether neighboring Pol III tRNA promoters showed a similar coordination of transcription initiation as observed above between Pol II promoters. Overall, we observed that chromatin fibers overlapping neighboring tRNA promoters were preferentially co-bound by initiating Pol III footprints significantly more frequently than expected by chance. In addition, we observed a strong distance-dependent relationship in this effect, with paired tRNA promoters located within 2.5 kb of each other showing the strongest coordination in Pol III co-occupancy **(Fig. 5d)**. In addition to having coordinated Pol III co-occupancy, tRNA genes located within 2.5 kb of each other also have significantly higher overall occupancy by Pol III transcription associated footprints, consistent with a model in which tRNA gene clustering enables higher overall expression via the spatial clustering of Pol III machinery **(Fig. 5e)**.

We next wanted to determine whether the Pol III coordination observed at tRNA genes was unique to tRNA genes or a general feature of Pol III-mediated transcription initiation. RNA polymerase III is also responsible for the transcription of 5S rRNA genes, which are 120 bp genes spatially clustered together as an array of ∼100 tandem copies of the same ∼375 bp repeat^62^. Of note, this tandem duplication is highly repetitive, making reference-based assessments of co-occupancy using short read sequencing approaches largely impossible. At 5S rRNA genes we found a similar pattern of footprints as at tRNA genes, corresponding to the selective occupancy at binding sites for TFIIIA, TFIIIB, and TFIIIC **(Fig. S10a,b)**. As individual Fiber-seq reads on average span ∼30 repeats of the 5S gene, we next determined whether Pol III 5S rRNA promoters showed a similar coordination of transcription initiation as observed above between Pol III tRNA promoters and between Pol II promoters. Overall, we observed that chromatin fibers overlapping neighboring 5S rRNA promoters were preferentially co-bound by Pol III transcription-associated footprints significantly more frequently than expected. In addition, we observed a strong distance-dependent relationship in this effect. However, this effect was longer range than that observed at neighboring tRNA promoters, with 5S rRNA genes showing coordinated occupancy by Pol III transcription associated footprints up to 10 copies away **(Fig. 5e)**. These results demonstrate that clustered, coordinated occupancy by Pol III is a consistent effect across the major Pol III transcribed genes, and further indicates that the condensed spatial clustering of 5S rRNA genes likely potentiates this effect.

### Transcription initiation is anti-coordinated between Pol II and Pol III genes

tRNA genes are often positioned adjacent to Pol II transcribed genes and the presence of these tRNA genes has been shown to disrupt the transcriptional output of these neighboring Pol II transcribed genes^63,64^. However, the exact mechanism driving the feedback between Pol II and Pol III transcribed genes is unknown. Current models suggest that tRNA genes may act as boundary elements in a Pol III transcription dependent manner^64^. To determine if transcription initiation at Pol III-transcribed tRNA genes influences Pol II-mediated transcription initiation at neighboring genes, we evaluated for transcription initiation footprints along chromatin fibers overlapping neighboring tRNA and Pol II-transcribed promoters. Overall, we observed that Pol III and Pol II genes showed the opposite effect as the coordination observed between pairs of Pol II promoters or pairs of Pol III genes. Specifically, we observed that PPP and PIC footprints were significantly depleted along chromatin fibers that contained Pol III transcription-associated footprints, and vice versa. This effect demonstrated a strong distance-dependent relationship, with anti-correlated Pol II and Pol III transcription initiation being observed for genes within 5 kb of each other **(Fig. 5g)**. Furthermore, as Pol III transcription-associated footprints at tRNA genes are significantly more prevalent than PPP and PIC footprints at Pol II-transcribed genes, the outcome largely results in Pol III mediated silencing of transcription at Pol II genes. Overall, these results demonstrate that tRNA mediated silencing of Pol II-transcribed genes is largely occurring at the level of transcription initiation.

## Discussion

Overall, we delineate at single-molecule and near single-nucleotide resolution both competition and coordination of transcription initiation and pausing and the surrounding nucleosome landscape along individual chromatin fibers. Specifically, we demonstrate both a mechanism for pause-inhibited initiation and pause-driven changes to the positioning and stability of nucleosome arrays that compete for the same genomic real estate. In addition, we demonstrate pervasive coordination of transcription initiation at nearby pairs of promoters and enhancer-promoter pairs, and anti-coordination between Pol II and Pol III transcribed genes. Together these findings illustrate transcription initiation as not merely a passenger occupying templated chromatin environments, but as an active participant in the regulation of gene expression and formation of chromatin architectures within these environments.

We find that Pol II pausing is directly responsible for a broad destabilization of the downstream nucleosome landscape up to 3 nucleosomes away from the TSS but has no effect on upstream nucleosomes. Furthermore, chromatin fibers maintain a short-term memory of these destabilized downstream nucleosomes at genes with frequent pausing. Upon pause release, fibers with destabilized downstream nucleosomes appear to be actively transcribed at a higher frequency, suggesting a direct connection between pausing, altered downstream nucleosome architecture, and elongation activity. Further, the pause-associated changes to promoter structure likely permit a more efficient reestablishment of both initiation and pausing upon pause release, via maintaining both an accessible TSS and pause site, which would otherwise be occluded by nucleosomes. Consequently, this directional, pause-specific, and expression-independent short-term memory may serve to prime highly paused genes for rapid, regulated transcription, consistent with the patterns of expression seen at developmental and heat shock genes^1,65^.

We also demonstrate the pervasive coordination of transcription initiation between spatially proximal Pol II transcribed promoters and enhancers. Coordination appears to be an effect shared across RNA polymerases and promoters and is largely limited to elements located within 5 kb of each other. Spatial proximity along the genome as a driver of synchronized, coregulated transcription suggests a functional basis for both proximal enhancer RNA expression and the clustering of Pol II and Pol III genes along the *Drosophila* genome. In addition, spatial coordination helps explain the marked variability in expression we observe between otherwise identical Pol III transcribed tRNA genes, with higher or lower transcription activity based on their proximity to other tRNA genes.

Together, these findings delineate the intricate dance that polymerases and the rest of the chromatin-bound proteome perform along individual chromatin fibers within the cell. In addition, this work highlights Fiber-seq as a central tool for unifying prior microscopy-, structural-, and genomic-based assays of RNA polymerase – establishing a single-molecule framework of competition and coordination of polymerase complexes with the surrounding nuclear landscape along individual chromatin fibers within each cell.

## Supplemental Figures

**Figure S1.**
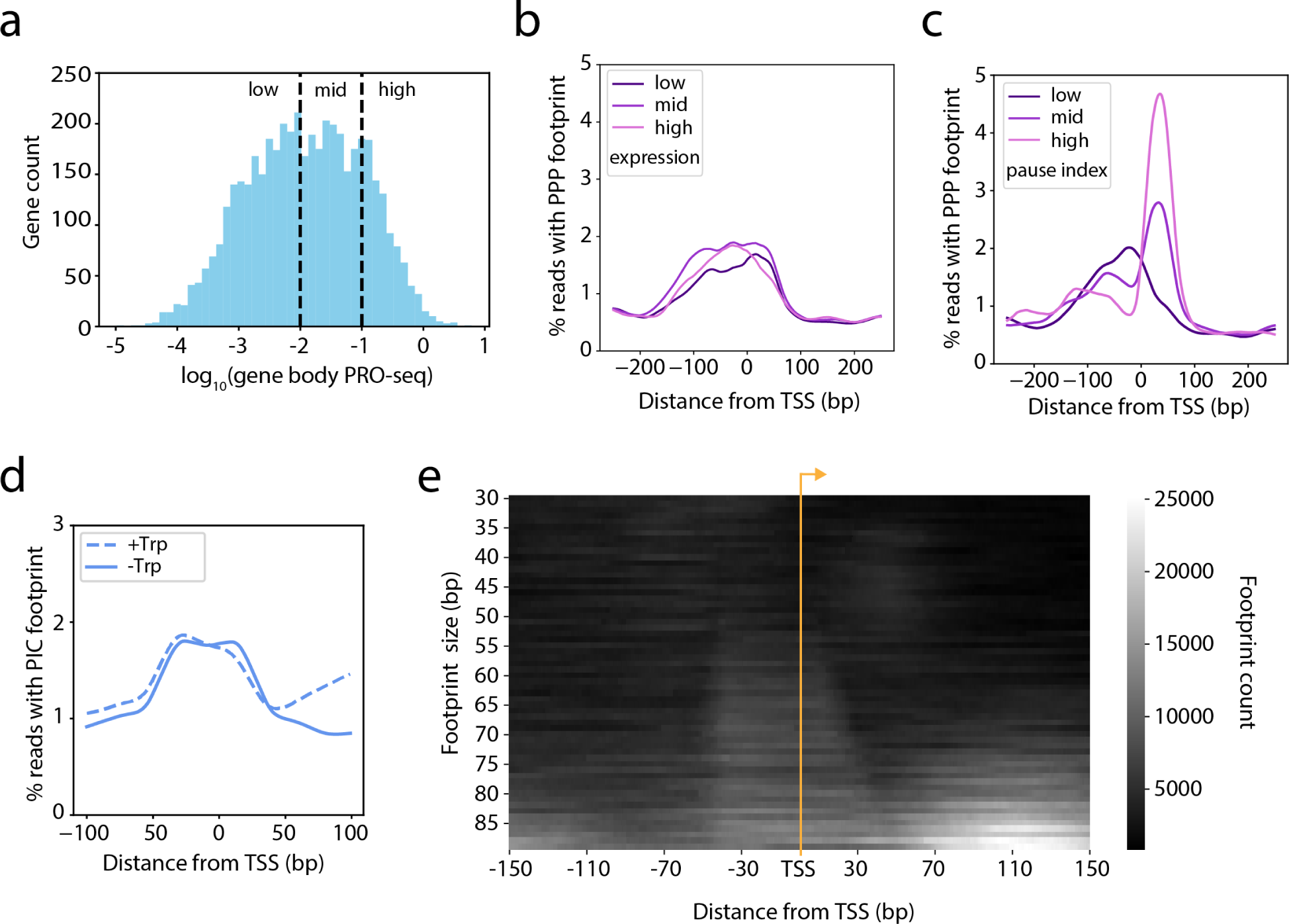
(related to Figure 1**) (a)** Histogram showing overall distribution and binning of expression (PRO-seq signal in gene body, normalized by gene length), as used throughout analyses. **(b)** Plot showing enrichment of PPP footprints at genes binned by expression level. **(c)** Plot showing enrichment of PPP footprints at genes binned by pause index. **(d)** Plot showing enrichment of PIC footprints with **(solid)** or without **(dashed)** triptolide. **(e)** Heatmap depicting the enrichment of differently sized Fiber-seq footprints with triptolide treatment at positions around transcription start sites (TSS) of genes with a pause index ≥10.

**Figure S2.**
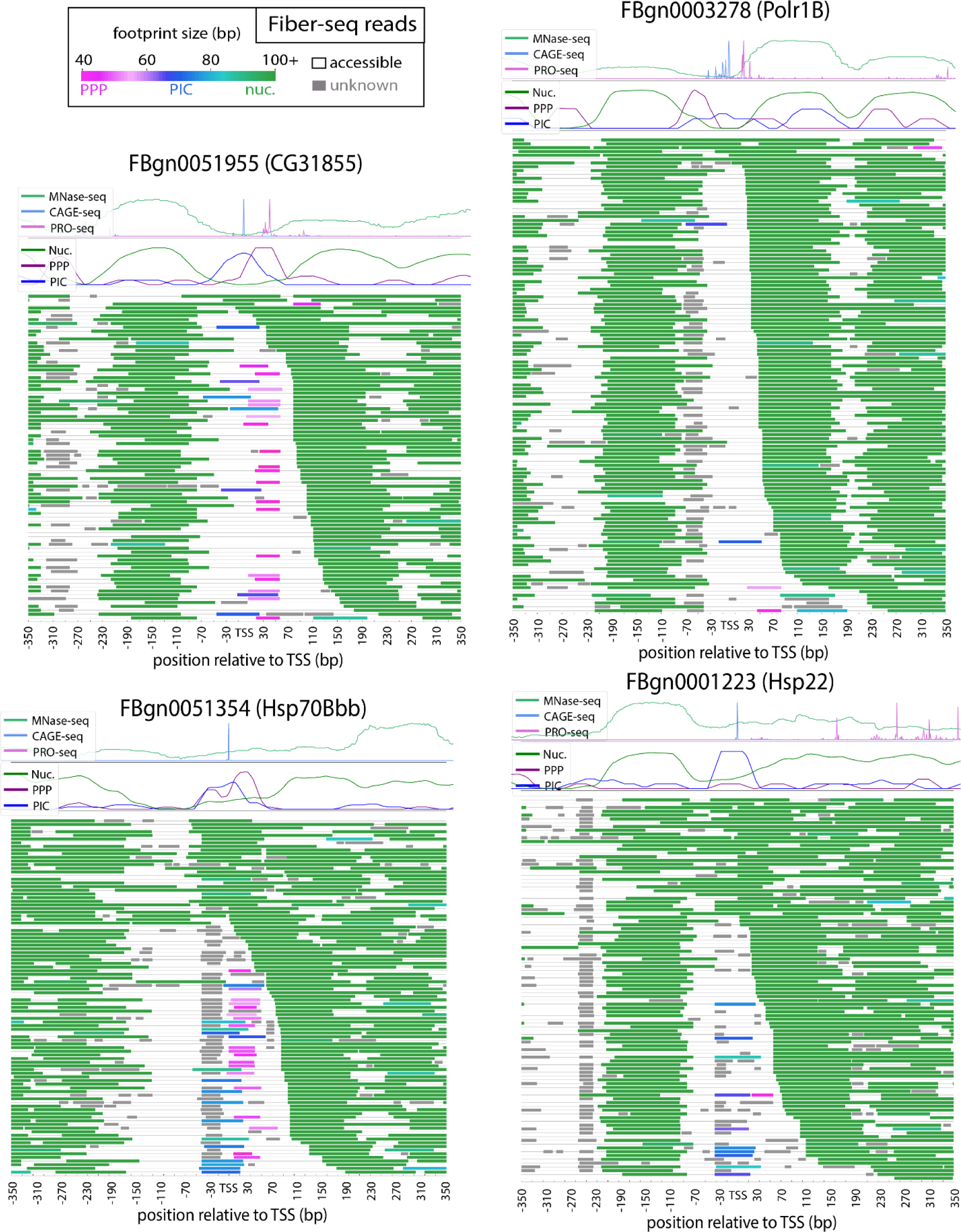
(related to Figure 1) Fiber-seq reads at four example protein coding loci. Each plot contains a track with MNase-seq, PRO-seq, and CAGE-seq, as well as a track showing enrichment of PPP, PIC, and nucleosome footprints for comparison. Below are all Fiber-seq reads aligned to each locus. Footprints are colored based on predicted identity (PPP = pink, PIC = blue, nucleosome = green, unknown = gray).

**Figure S3.**
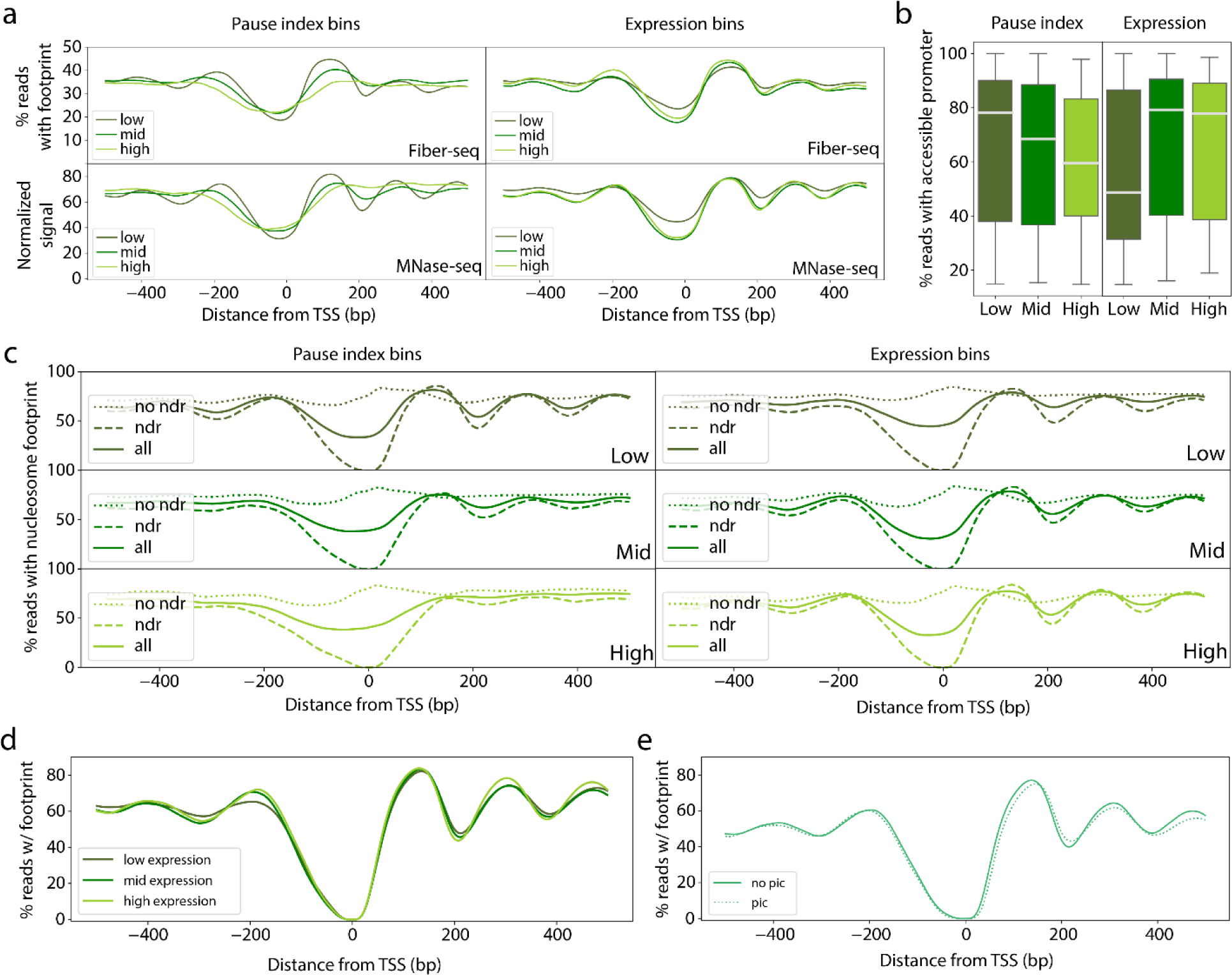
(related to Figure 2) **(a)** Comparison between **(top)** MNase-seq signal and **(bottom)** Fiber-seq nucleosome enrichment at **(left)** different pausing levels or **(right)** different expression levels. **(b)** Box plot showing the fraction of Fiber-seq reads with an accessible promoter at **(left)** different pausing levels or **(right)** different expression levels. **(c)** Comparison between Fiber-seq nucleosome enrichment at **(left)** different pausing levels or **(right)** different expression levels with **(dashed)** or without **(dotted)** an accessible promoter, in comparison to the nucleosome enrichment across all genes. **(d)** Enrichment of nucleosome footprints in Fiber-seq reads at different levels of expression, only including reads with an accessible promoter. **(e)** Comparison of nucleosome enrichment in Fiber-seq reads with **(dotted)** or without **(solid)** a PIC footprint, only including reads with an accessible promoter and sampled to capture an equal amount of PIC and no-PIC reads from each gene.

**Figure S4.**
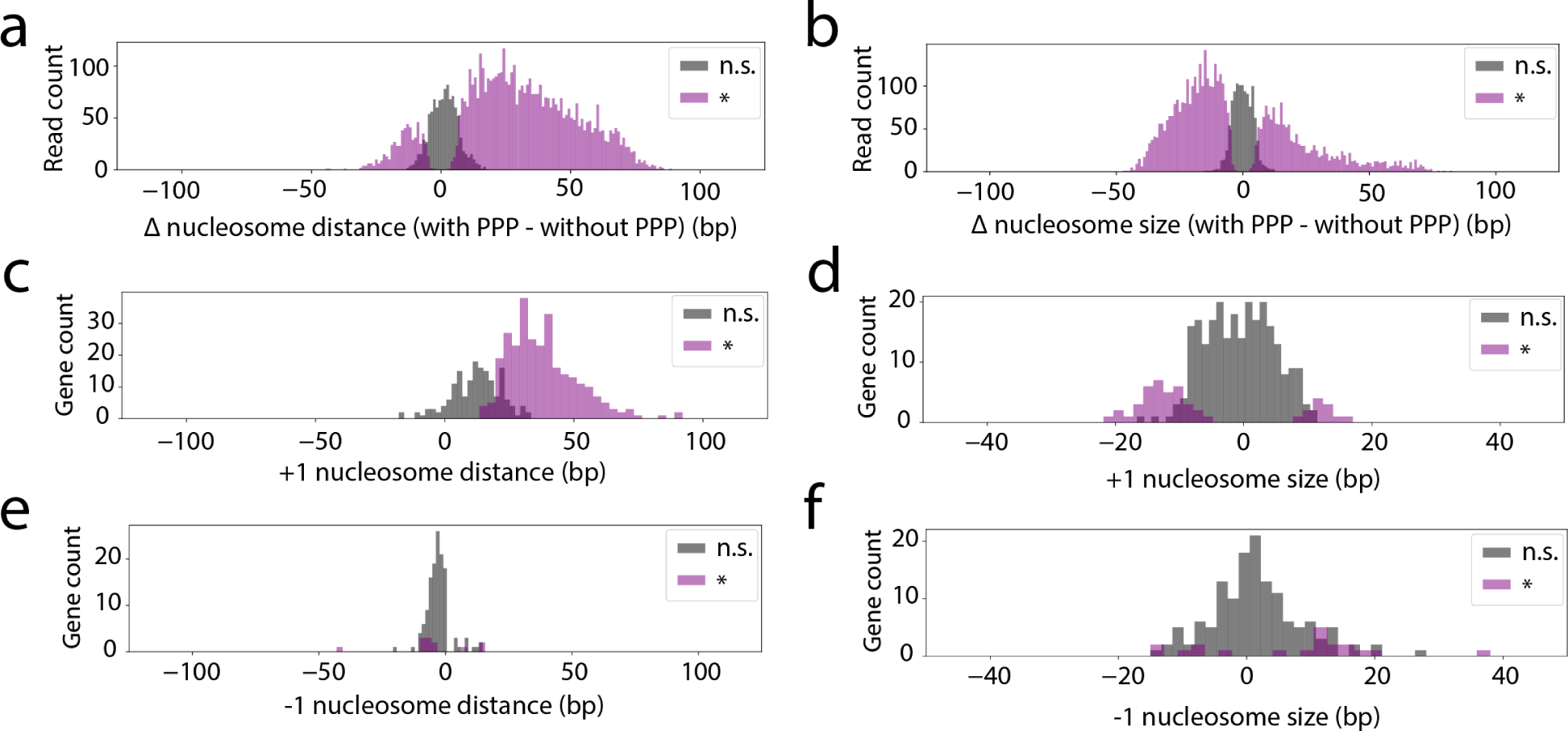
(related to Figure 2) **(a,b)** Each individual read with a PPP footprint was compared to the bootstrapped mean distance **(a)** and size **(b)** of the +1 nucleosome. Histogram of difference in +1 distance or size for each PPP read compared to the bootstrapped mean +1 distance or size. Bins are colored and divided based on their associated −log_10_(p-value), generated by comparing the value to the bootstrapped distribution at the origin locus (empirical confidence interval from bootstrapped mean, n.s. signifies p > .05, * signifies p-value < .05). **(c-f)** Comparing reads with or without a PPP footprint at each locus with at least 5 PPP reads. Graphs in order are **(c)** +1 distance, **(d)** +1 size, **(e)** −1 distance, **(f)** −1 size. Histogram of the mean distance or size of the +1 or −1 nucleosome for reads with a PPP footprint at each locus, shaded based on significance (Wilcoxon Rank Sum test, n.s. signifies p > .1, * signifies p-value < .1).

**Figure S5.**
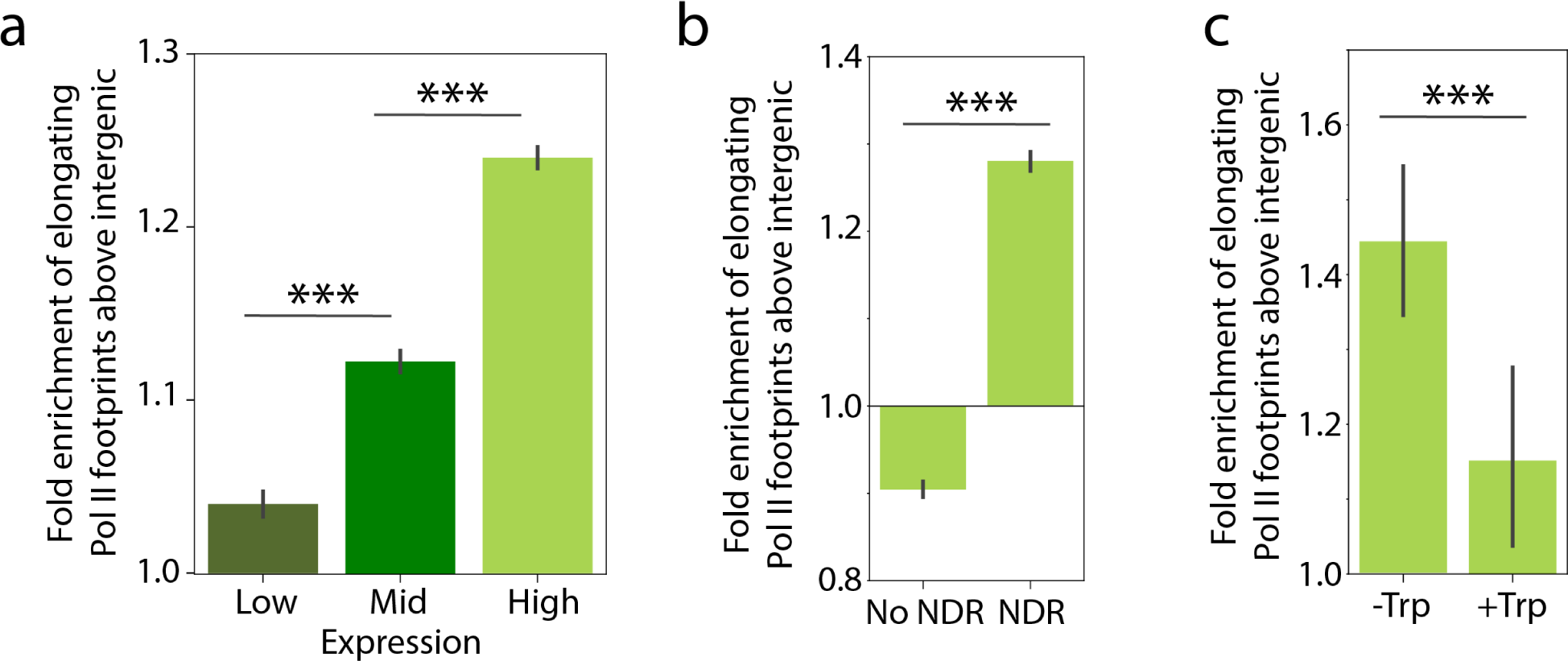
(related to Figure 2) **(a)** Bar plot quantifying the fold enrichment of putative elongating Pol II footprints within gene bodies of genes split by expression based on PRO-seq signal within gene bodies Expression values for genes are binned into “high” (≥100), “mid” (≥10, <100), and “low” (<10) values. Pairwise significance is indicated above (two-sample t-test, *** signifies p-value < 0.001). **(b)** Bar plot quantifying the fold enrichment of putative elongating Pol II footprints within gene bodies at reads with an inaccessible or accessible promoter at the corresponding gene. Pairwise significance is indicated above (two-sample t-test, *** signifies p-value < 0.001). **(c)** Bar plot quantifying the fold enrichment of putative elongating Pol II footprints within gene bodies at reads with a PIC footprint at the corresponding gene. Pairwise significance between the −Trp and +Trp datasets is indicated above (two-sample t-test, *** signifies p-value < 0.001).

**Figure S6.**
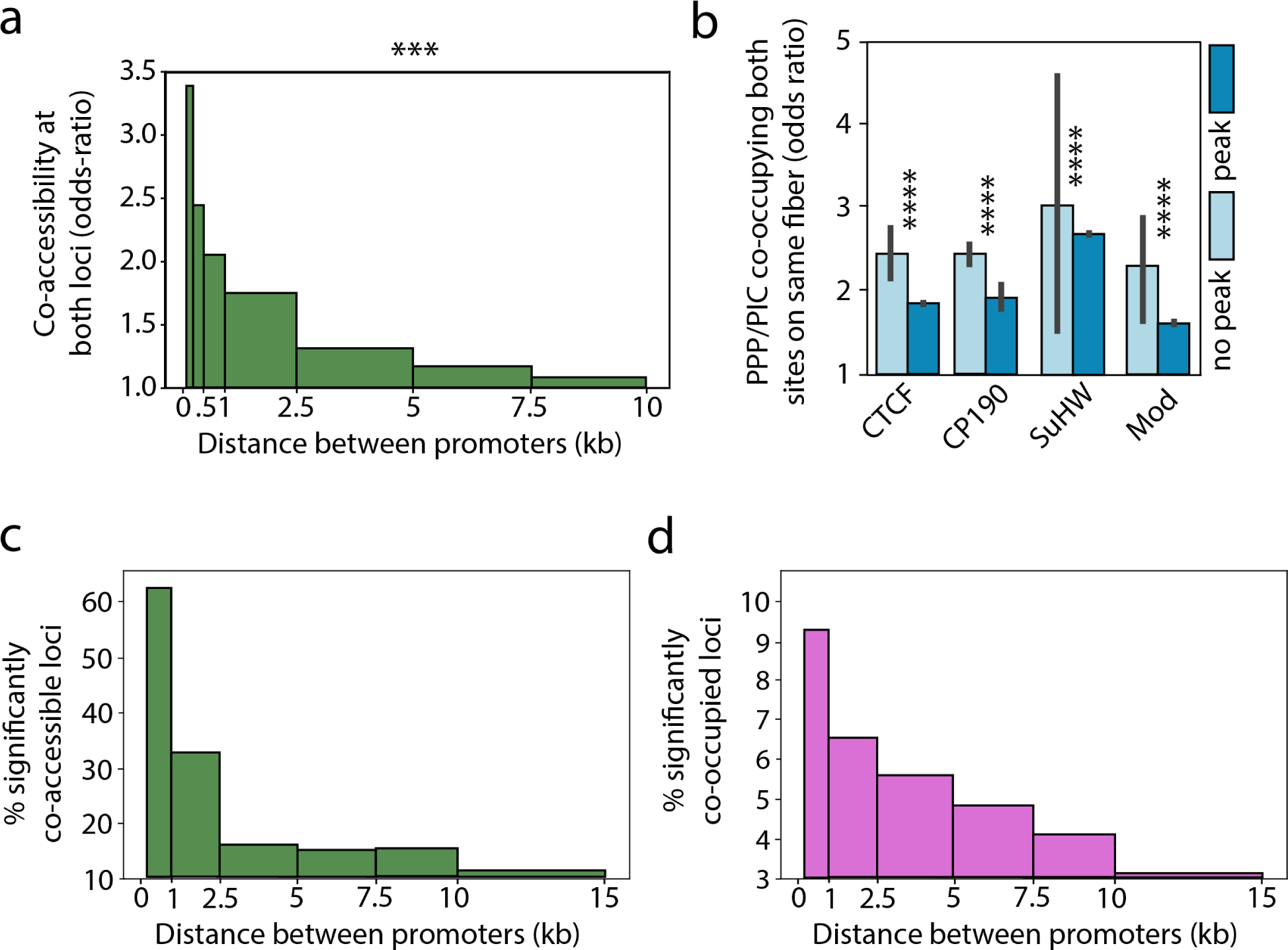
(related to Figure 4) **(a)** Bar plot showing the Fisher’s exact test odds ratio of co-accessibility of promoters on reads overlapping promoter pairs binned by distance. Significance is schematized above the plot (Fisher’s exact test, *** signifies p-value < 0.001). **(b)** Bar plot depicting the coordination of transcription initiation pairs of genes separated or not separated by a ChIP-seq peak of individual insulator binding proteins included in the pooled analysis from Figure 4. All pairs are found within 2.5kb of each other and reads from each pair of groups are sampled to capture an identical count of reads from pairs at any given distance, with error bars corresponding to the confidence interval calculated from 10,000x sampling iterations. Pairwise significance is indicated above (two-sample t-test, **** signifies p-value < 10^−200^). **(c)** Bar plot showing the percentage of pairs of genes with a significant (p-value < 0.1) Fisher’s exact test odds ratio of co-accessibility. **(d)** Bar plot showing the percentage of pairs of genes with a significant (p-value < 0.1) Fisher’s exact test odds ratio of co-occupancy by PPP or PIC footprints.

**Figure S7.**
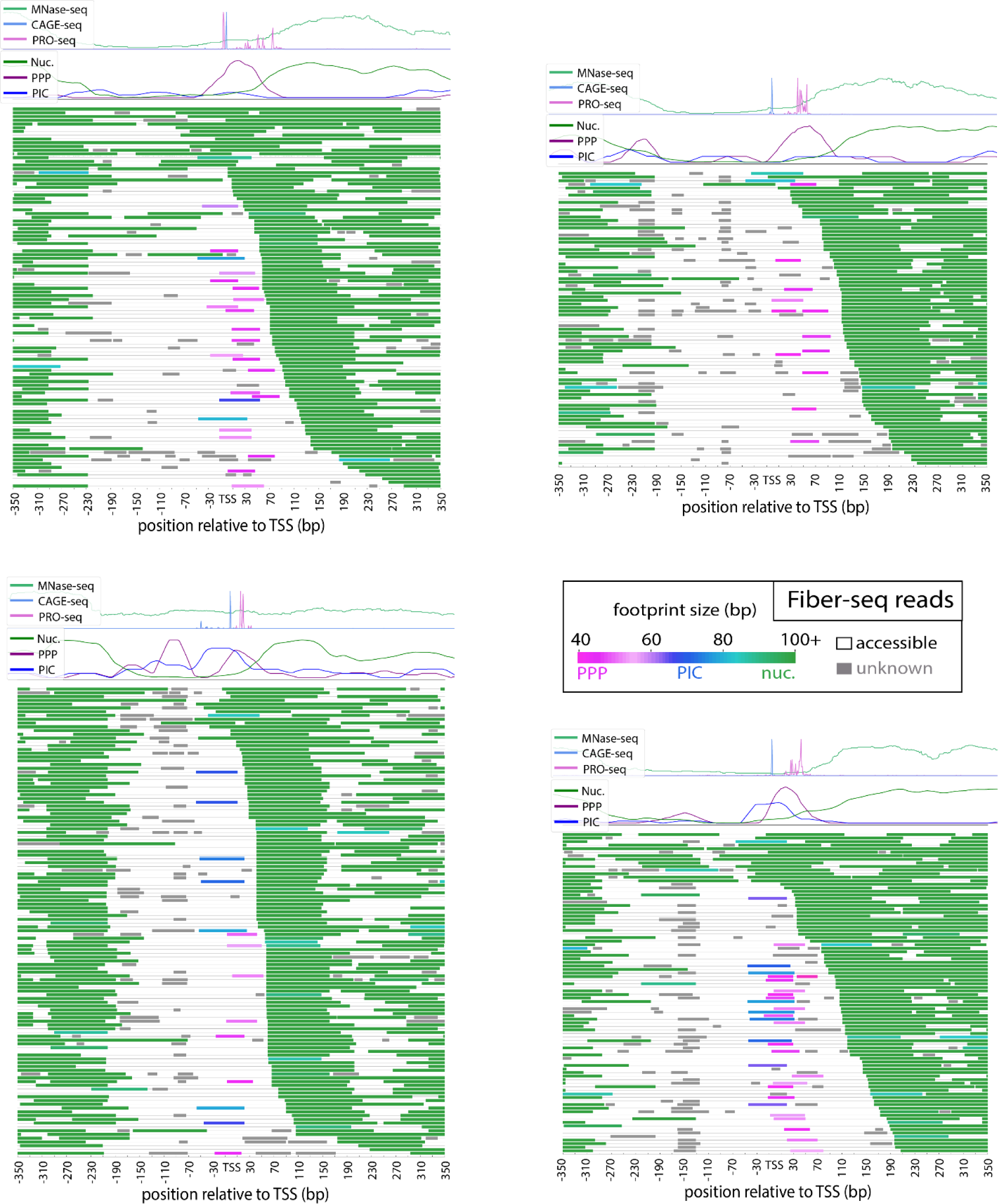
(related to Figure 4) Fiber-seq reads at four example transcribed enhancer loci. Each plot contains a track with MNase-seq, PRO-seq, and CAGE-seq, as well as a track showing enrichment of PPP, PIC, and nucleosome footprints for comparison. Below are all Fiber-seq reads aligned to each locus. Footprints are colored based on predicted identity (PPP = pink, PIC = blue, nucleosome = green, unknown = gray).

**Figure S8.**
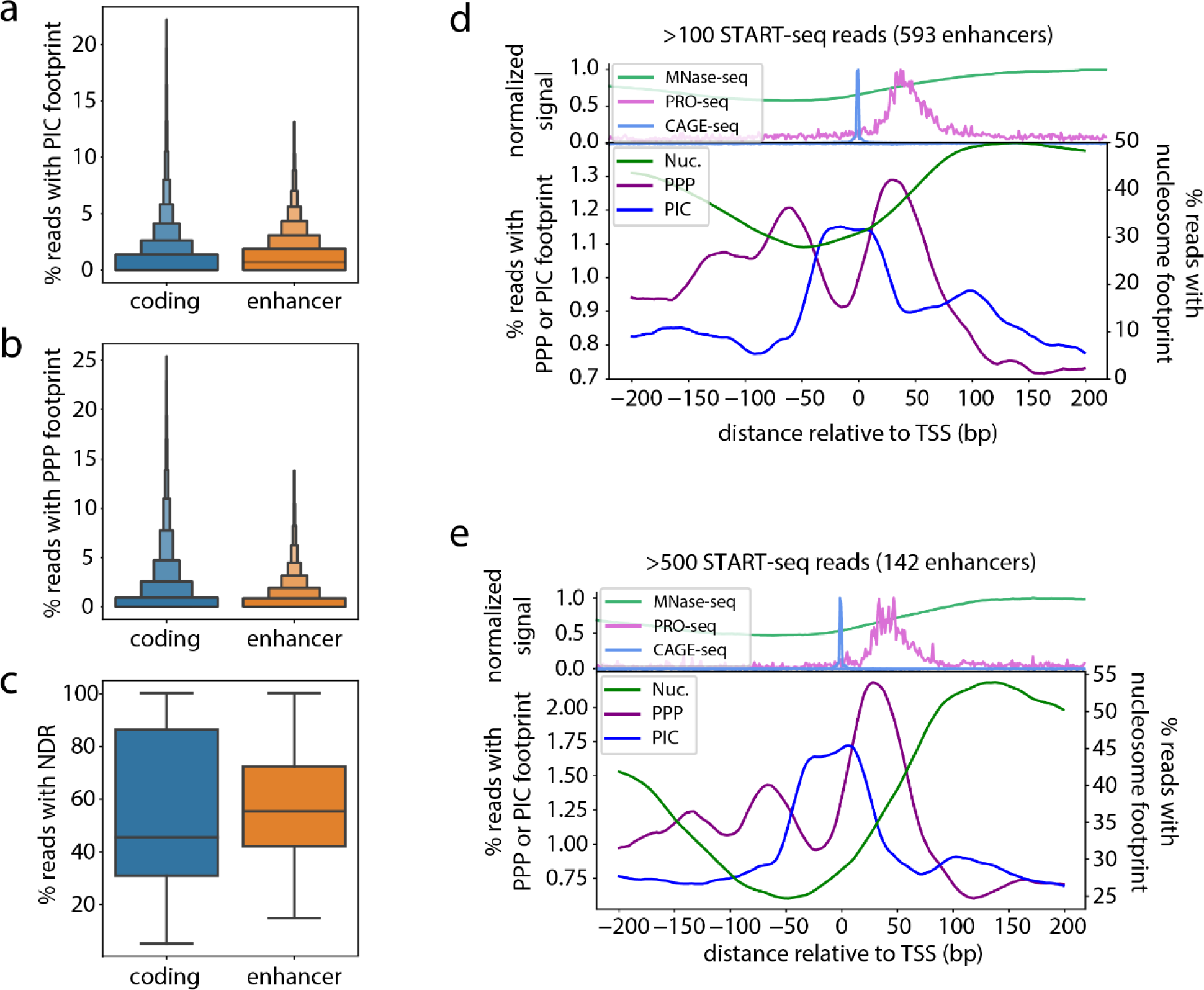
(related to Figure 4) **(a)** Boxenplot showing the distribution of percentage of reads with PIC footprints at protein-coding genes and transcribed enhancers. Pairwise significance is indicated above (two-sample t-test, *** signifies p-value < 0.001). **(b)** Boxenplot showing the distribution of percentage of reads with PPP footprints at protein-coding genes and transcribed enhancers. Pairwise significance is indicated above (two-sample t-test, *** signifies p-value < 0.001). **(c)** Boxplot showing the percentage of reads with an accessible promoter for protein coding genes and transcribed enhancers. Pairwise significance is indicated above (two-sample t-test, *** signifies p-value < 0.001). **(d,e)** Plots showing **(top)** MNase-seq, PRO-seq, and START-seq signal at transcribed enhancers compared to **(bottom)** enrichment of PPP, PIC and nucleosome footprints in Fiber-seq reads. Each plot has the indicated (50, 100) minimum coverage of START-seq for the genes included in the plot.

**Figure S9.**
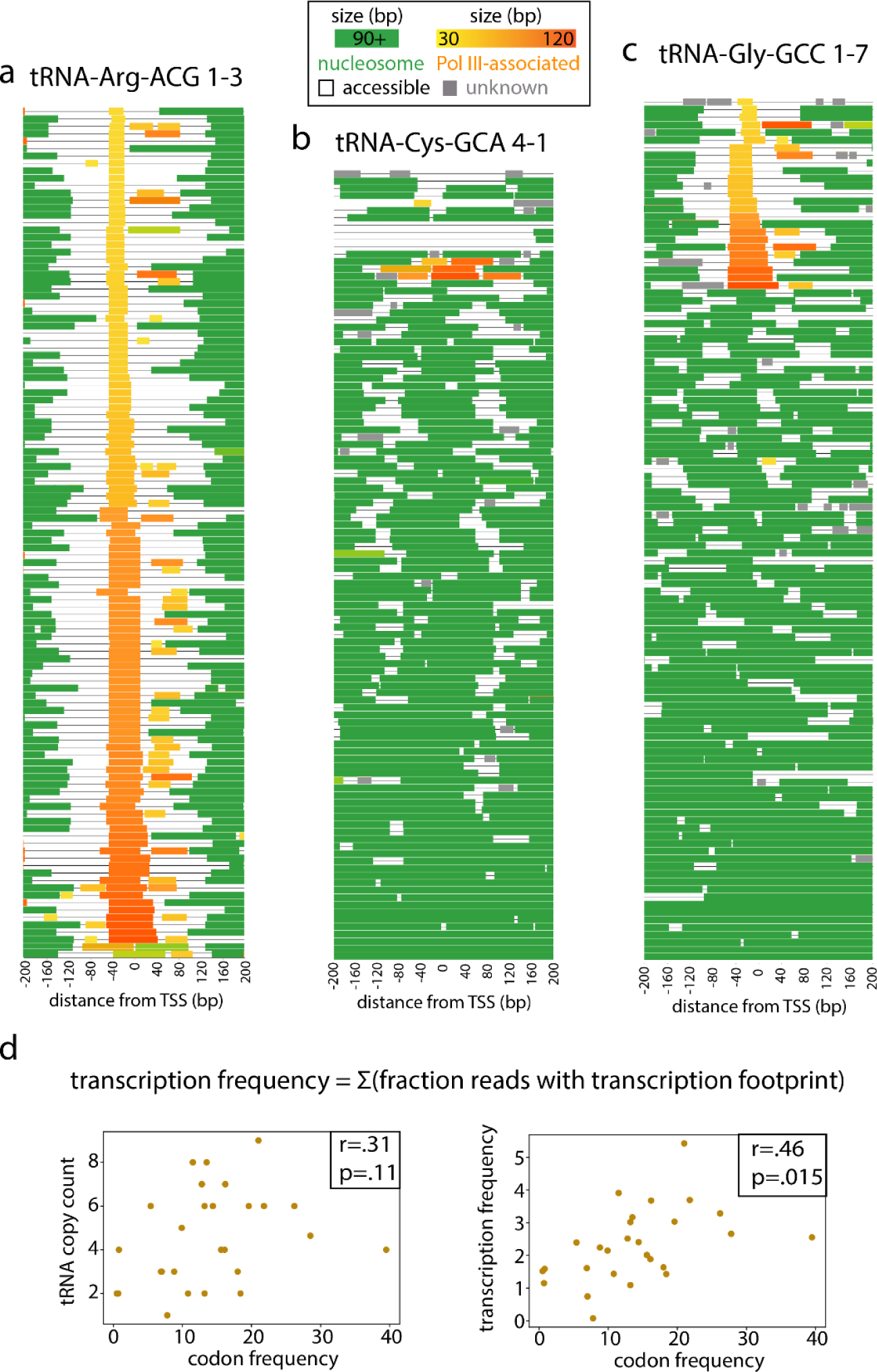
(related to Figure 5) **(a,b,c)** Fiber-seq reads at three example tRNA loci. Footprints are colored based on predicted identity (nucleosome = green, tRNA transcription associated footprints = yellow to orange based on size). **(d)** Transcription frequency scores were calculated for each family of tRNAs with identical anticodon sequences and with all members having coverage in the middle 95% of the distribution of overall Fiber-seq sequencing coverage. **(left)** Scatter plot depicting tRNA family gene copy number plotted against their corresponding codon frequency in the *D. melanogaster* genome. **(right)** Scatterplot depicting tRNA isodecoder transcription frequency plotted against corresponding codon frequency in the *D. melanogaster* genome. The Pearson correlation and significance are shown in the top right corner of each plot.

**Figure S10.**
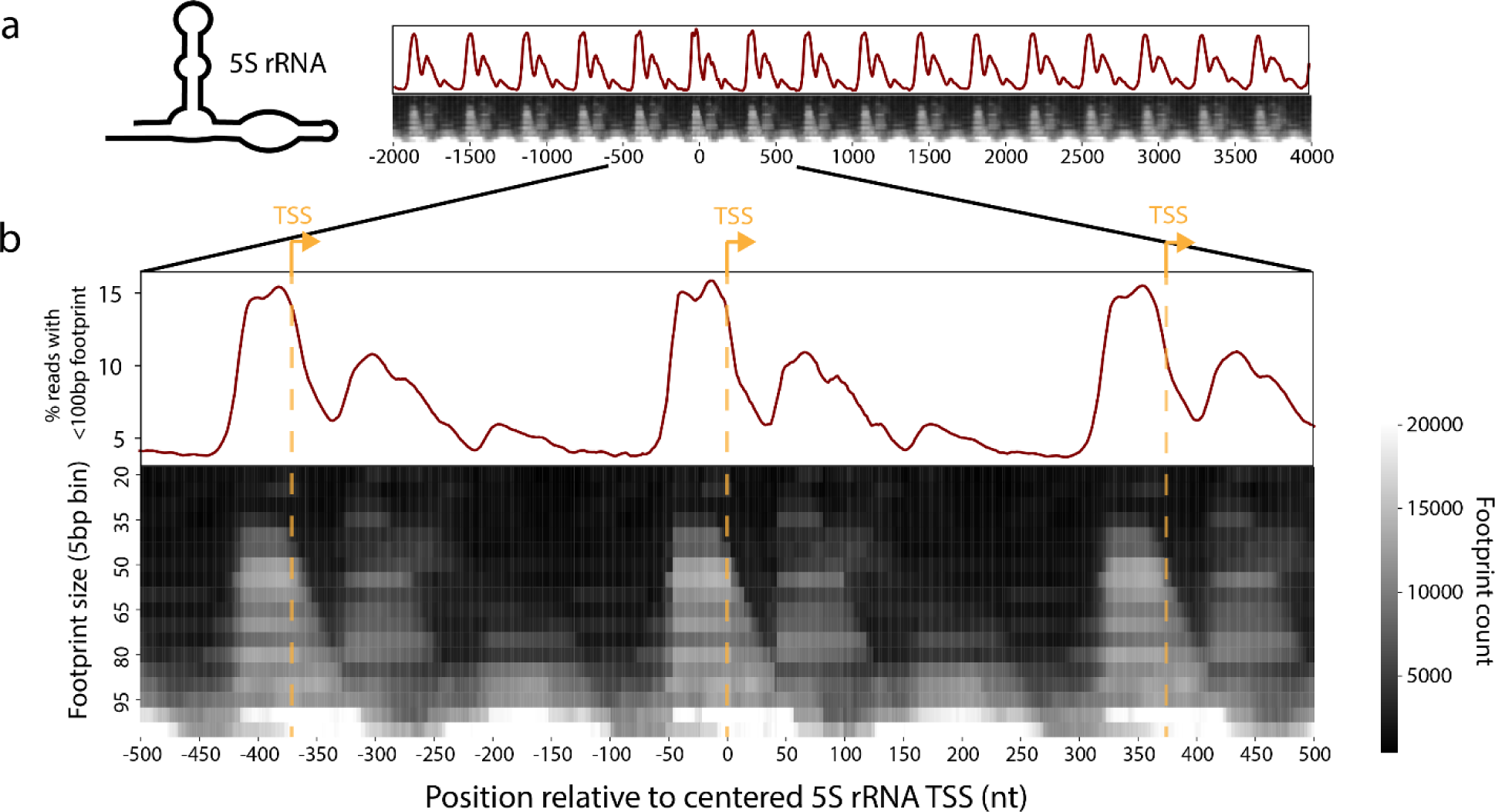
(related to Figure 5) **(a)** Heatmaps showing enrichment of differently sized footprints at 5S rRNA genes. **(top)** Wide view showing 16? 5S rRNA copies. **(bottom)** Zoomed in view showing 3 5S rRNA genes. For both wide and zoomed plots: **(top)** enrichment of 30-90 bp footprints relative to 5S rRNA TSSs and (**bottom**) heatmap depicting the enrichment of differently sized Fiber-seq footprints with respect to 5S rRNA TSSs.

## Supplemental Tables

**Table S1–.**
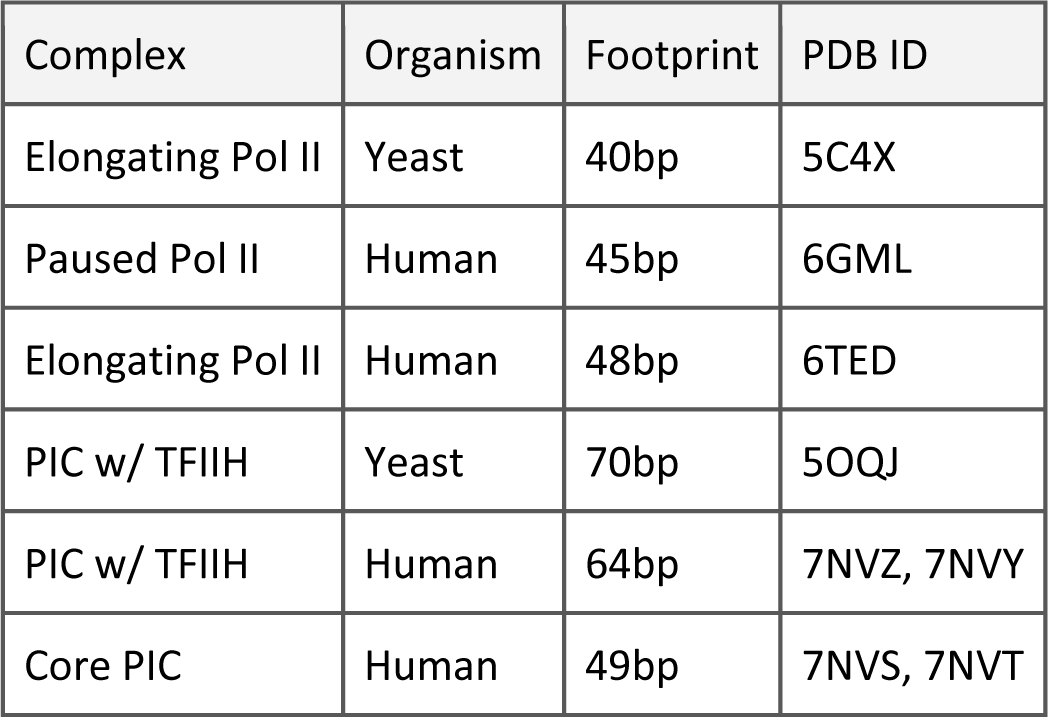
footprint sizes compared to structures/ChIP.

## Methods

### Datasets

Previously published datasets used in our analyses include: processed PRO-seq (GSE42117), MNase-seq datasets (GSE128689), CAGE-seq datasets (ENCSR863NJP), TAD boundaries (GSE58821), ChIP-seq peaks for insulator binding proteins (GSE41354), START-seq data (GSE85191), enhancer locations (GSE57876). Codon usage was based on the Codon Usage Tabulated from GenBank database^61^. All datasets originally aligned to the Drosophila dm3 assembly were lifted over to the dm6 annotation using the UCSC liftover utility.

### Defining CAGE-seq, START-seq and PRO-seq peaks

To call CAGE- and PRO-seq peaks near TSSs, a sliding 20 bp window from −50 to +50 bp relative to the TSS (CAGE-seq) or 0 to 100 bp relative to the TSS (PRO-seq) around the TSS was used, assigning maxima within each window as peaks. A minimum of 10 reads was required to call a peak, with the strongest peak above that threshold being assigned as the primary PRO-seq or CAGE-seq peak. All secondary peaks were also required to have a minimum of 10-fold lower coverage than the primary peak. In the case of the primary CAGE-seq peak, the TSS used in subsequent analyses was adjusted from the reference annotation to that peak if the reference differed.

### Calculating pause index and expression levels

Pause index and expression were both calculated using PRO-seq data. Pause index was calculated as the ratio of PRO-seq coverage in the promoter region (−100 to 300 bp relative to the TSS) to the body of the gene (300 bp after the TSS to the end of the gene), normalized by gene length. Expression was calculated based on PRO-seq coverage in the gene body (300 bp after the TSS to the end of the gene) normalized by gene length.

### Hia5 expression and purification

pHia5ET was expressed and purified as previously described^22^. pHia5ET was transformed into T7 Express lysY/Iq Escherichia coli cells (NEB C3013I). Overnight cultures were added to two 1 L cultures of LB medium supplemented with 50 µg/mL kanamycin and grown with shaking at 37°C to an OD600 of 0.8-1.0. Isopropyl β-D-1-thiogalactopyranoside (IPTG) was added to a final concentration of 1 mM and protein was expressed for 4 hours at 20°C with shaking. Cells were pelleted at 5,000 rpm for 10 minutes at 4°C. The pellet was resuspended in 35 mL lysis buffer (50 mM HEPES, pH 7.5; 300 mM NaCl; 10% glycerol; 0.5% Triton X-100; 10 mM β-mercaptoethanol) supplemented with 2X Complete, EDTA-free Protease Inhibitor Cocktail (Millipore Sigma 11873580001). Cells were lysed by probe sonication (Qsonica Q125) for 10 minutes on ice at 50% amplitude, 30 seconds on/off. Lysate was clarified by centrifuging for 1 hour at 40,000 x g. Ni-NTA Agarose (Qiagen 30210) was prepared by washing 5 mL slurry with 30 mL equilibration buffer (50 mM HEPES, pH 7.5; 300 mM NaCl; 20 mM Imidazole) and centrifuged at 500 x g for 3 minutes, repeating once. The clarified lysate and Ni-NTA agarose were combined and rotated at 4°C for 1 hour. The lysate mixture was poured over a disposable gravity flow column (Bio-Rad 7321010) and washed with 20 mL Buffer 1 (50 mM HEPES, pH 7.5; 300 mM NaCl; 50 mM imidazole) and 15 mL Buffer 2 (50 mM HEPES, pH 7.5; 300 mM NaCl; 70 mM imidazole). Protein was eluted with 15 mL Elution Buffer (50 mM HEPES, pH 7.5; 300 mM NaCl; 250 mM imidazole). 6-8K MWCO SnakeSkin dialysis tubing (Spectrum Laboratories 132650) was pre-wet in dialysis buffer (50 mM Tris-HCl, pH 8; 100 mM NaCl; 1 mM DTT) and the eluate was added to the tubing and dialyzed against 2 L dialysis buffer overnight at 4°C. Dialyzed sample was concentrated using a 10K Amicon Ultra-15 spin concentrator (Millipore Sigma UFC901008) and centrifuged at 3,220 x g to concentrate to a volume of less than 1 mL. The concentrated sample was injected onto Tandem HiTrap Q HP (Cytiva 17115301) and HiTrap SP HP (Cytiva 17115101) columns equilibrated with FPLC Buffer A (50 mM Tris-HCl, pH 8; 100 mM NaCl; 1 mM DTT). Columns were washed with 5 column volumes of FPLC Buffer A. The Q column was removed and the sample eluted from the SP column using a linear gradient over 20 column volumes of 0 to 100% FPLC Buffer B (50 mM Tris-HCl, pH 8; 1 M NaCl; 1 mM DTT). Peak fractions were collected and concentrated to 250 µL using a 10K Amicon Ultra-15 spin concentrator. Protein was supplemented with glycerol to a final concentration of 10%, frozen, and stored at −80°C. Protein purity was assessed by SDS-PAGE, and its activity measured by an *in vitro* methylation assay.

### Fiber-seq

Fiber-seq was carried out as previously described^22^ on *D. melanogaster* S2 cells. Cells were grown in the dark in 75 cm^2^ flasks to a final density of 3 million cells/mL, with 8 million cells used per sample. Cells were then spun down at 250 x g for 5 minutes, resuspended in 1 mL of 1x DPBS, spun down again at 250 x g for 5 minutes, and resuspended in 150 µL Buffer A (15 mM Tris-Cl pH 8.0, 15 mM NaCl, 60mM KCl, 1 mM EDTA, 0.5 mM EGTA, 0.5 mM Spermidine). To isolate nuclei, a 0.05% final concentration of IGEPAL was added and cells were incubated for 10 minutes on ice and subsequently spun down at 500 x g for 5 minutes. The nuclei pellets were then resuspended in 60 µL Buffer A with the addition of 0.8 mM SAM and 200 U of Hia5, and incubated for 10 minutes at 25°C. The reaction was quenched with the addition of 1% final SDS and mixing using a wide-bore pipette tip. DNA was then purified using the Promega Wizard HMW DNA Extraction Kit (Promega A2920) and submitted for library preparation and sequencing.

Four sets of biological replicates were generated in S2 cells with no perturbations, with the latter 3 split into two technical replicates each -- a total of seven datasets overall. Two biological replicates in S2 cells treated with triptolide were also generated, with the latter biological replicate split into two technical replicates. For the samples treated with triptolide, a final concentration of 10 μM of triptolide was added to the cells 30 minutes prior to harvesting and an identical molarity maintained through the Fiber-seq protocol. Two control Fiber-seq datasets were generated as inputs for FiberHMM. The control Fiber-seq datasets were carried out identically, except that the no-Hia5 sample had no added Hia5 and that the dechromatinized sample was carried out on purified genomic *Drosophila* S2 genomic DNA instead. Specifically, control Fiber-seq on dechromatinized DNA was performed on genomic DNA purified from *Drosophila* S2 cells using the Promega Wizard HMW DNA Extraction Kit (Promega A2920). 2 µg of DNA was used in a final volume of 60 µL in Buffer A with 0.8 mM S-adenosylmethionine and 200 U Hia5. Samples were incubated for 2 hours at 37°C before quenching by purification with the Promega Wizard HMW DNA extraction kit.

### Fiber-seq processing

Fiber-seq datasets were assigned circular consensus sequences using the CCS tool (6.4.0) from Pacific Biosciences. They were then mapped to the dm6 genome using the pbmm2 package. Methylations were then called using the fibertools package (0.3.0) in SCNN mode and reads were converted to a BED file using a threshold of 250 for the m6A score.

### FiberHMM

Footprints were called on Fiber-seq reads using FiberHMM. FiberHMM is based on a hidden Markov model (HMM) with two hidden states--accessible and inaccessible. To account for sequence-related biases from Hia5 or the methylation caller, at each position the model takes into account both the base and its surrounding 6 nt sequence context. The emission probabilities used in the model are the probabilities of methylation of a given base with its +/− 3 nt sequence context in an accessible or inaccessible state based on experimental-derived methylation rates from control datasets. The probability of methylation given an accessible state was based on the methylation frequency in a dataset generated from dechromatinized Drosophila S2 cell genomic DNA. The probability of methylation given an inaccessible state was based on the methylation frequency in a dataset generated from Drosophila S2 cells untreated with Hia5. Transition and starting probabilities for the HMM were trained 20 times on 1000 reads sampled in equal proportions from all untreated datasets and removed from subsequent analyses, with initial probabilities picked from the Dirichlet distribution with all parameters set to 1. The best model was chosen and then used for all subsequent footprint calling.

### Identifying size ranges for nucleosome footprints

Nucleosomes in general were defined as footprints greater than 90 bp. Most nucleosome footprints are single nucleosomes, which range from 90-200 bp, which is the definition used for analyses involving positioning and distance of single nucleosomes, specifically those featured in Figure 2.

### Identifying size ranges for PPP and PIC footprints

To identify the size ranges of PPP and PIC footprints, Fiber-seq reads overlapping promoters with a pause index greater than 10 were identified. These reads were then aligned around the primary TSS of each promoter, plotting the enrichment of different footprint sizes at positions around the TSS in the form of a heatmap (Fig. 1c). This heatmap was then used to identify size ranges of two enriched footprint populations around the TSS corresponding to putative PPP and PIC footprints--40-60 bp and 60-80 bp respectively.

### Defining PPP and PIC footprints

PPP and PIC footprints were defined using two requirements. The first requirement was that the footprint must be within the expected size range, and the second was that the footprint must have a PRO-seq or CAGE-seq peak overlapping the middle 70% of the footprint and that the PRO-seq peak or CAGE-seq peak be within a range of 0 to 100 bp or +/− 50 bp around the TSS respectively. Due to higher false positive rates of PPP calls at genes with a very low pause index, a slightly stricter definition of the PPP footprint was used for the analysis in Figure 2, requiring that the PRO-seq peak overlap the middle 50% of the putative PPP footprint.

### Identifying reads with an accessible promoter

Reads with an accessible promoter for a given gene were defined as those reads lacking a nucleosome footprint overlapping within +/−10 bp around the primary TSS of that gene.

### Validation of footprint sizes from existing structures

To compare footprint sizes to structures, structures from the referenced studies were accessed and downloaded from RCSB PDB (accession codes: 5C4X, 6GML, 6TED, 5OQJ, 7NVZ, 7NVS, 7NVT, and 7NVY) and visualized in PyMOL. The scaffold DNA bases obscured by the protein structure were counted manually to estimate the predicted footprint size associated with the structure.

### Sampling reads based on PPP and PIC footprints

To allow for direct comparison between features of reads with or without a PPP or PIC footprint on a global scale a sampling approach was used. At each gene all reads with an accessible promoter were identified. These reads were then split based on if they contained a PPP footprint, a PIC footprint, or neither. Reads at each gene were then sampled to match the count of reads with a PPP footprint to those with neither a PPP or PIC, or to match the count of reads with a PIC footprints to the count of those with neither a PPP or PIC. All analyses involving sampling were repeated with a different randomization seed at least 50 times to make sure that any conclusions were stable and not a result of sampling biases.

### Sampling reads based on distance

For comparison of the co-occupancy of PPP or PIC footprints at pairs of genes with or without an interrupting TAD boundary or insulator binding protein ChIP-seq peak, a similar sampling method was carried out as described above. The distance between each possible pair of genes was calculated, and the pairs included in the analysis were sampled from each group to capture an equal distribution of distances for each condition (with or without a TAD boundary or insulator binding protein ChIP-seq peak). This sampling was repeated 10,000x to capture the sampling variance and generate a confidence interval.

### Segmenting downstream and upstream nucleosomes

To identify the most likely position of nucleosomes upstream and downstream of the −1 and +1 nucleosome respectively, a Gaussian mixture model (GMM) was used. The GMM was trained on the distribution of nucleosome footprints either upstream or downstream to identify 95% confidence intervals for each nucleosome within the range. The number of states for the GMM was chosen based on a visual approximation of the number of peaks from the overall distribution.

### +1 and −1 nucleosome size and distance on a per-gene basis

To compare nucleosome footprint distance and size on a per-gene basis **(Fig. S4c-f)**, the size and distance of the +1 or −1 nucleosome footprints were calculated across all reads at each individual locus and a Wilcoxon Rank Sum test was carried out, comparing the reads with a PPP footprint to those without a PPP footprint.

### +1 nucleosome size and distance on a per-read basis

To compare nucleosome footprint distance and size on a per-read basis **(Fig. S4a,b)**, the size and distance of the +1 nucleosome were calculated across all reads at each individual gene. These features were then bootstrap resampled across all reads containing an accessible promoter at each locus to calculate a mean and confidence interval for each feature at each locus. Each read with a PPP footprint was then tested against the corresponding mean and confidence interval to find the difference and significance of the +1 nucleosome footprint distance or size observed in that read compared to the other reads at the origin gene.

### Defining elongating Pol II footprints

Elongating Pol II footprints were defined based on matching the size range of a PPP footprint, 40-60 bp, and existing in gene bodies outside of a range of +/−300 bp around any annotated TSS or predicted eRNA TSS in the genome. The frequency of these footprints was defined as the count of the footprints per kilobase. The fold enrichment of elongating Pol II footprints was calculated based on the enrichment of identical footprints in intergenic regions outside of a range of +/−300bp from an annotated TSS or predicted eRNA TSS in the genome in the dataset from which that read originated.

### Pause-inhibited initiation quantification

To quantify the inhibition of simultaneous PPP and PIC occupancy, all genes which had at least one PPP and PIC footprint across all corresponding Fiber-seq reads were identified. These genes were then binned based on the distance between the primary CAGE-seq and PRO-seq peak and a contingency table was generated for the count of individual, simultaneous, and absent PPP and PIC footprints at these binned loci. Fisher’s exact test was then used to determine the odds ratio and p-value of having a simultaneous PPP and PIC footprint within each distance bin given the contingency table.

### Identifying transcribed enhancers

Transcribed enhancers were called by identifying TSSs within a 1kb window of enhancers from STARR-seq using START-seq data. Enhancer TSSs could not be within 500bp of any annotated TSS in the genome, including pseudogenes, noncoding RNAs, and other transcribed enhancers with higher maximum START-seq signal.

### Defining tRNA Pol III transcription-associated footprints

A similar strategy as the one employed to identify PPP and PIC footprints was used to identify Pol III transcription associated footprints. First, a heatmap was generated to identify regions and sizes of enriched footprints aligned around all tRNA TSSs. Several unique populations of footprints were visible, overlapping known binding sites of TFIIIB and TFIIIC, with sizes ranging from 30-140 bp. We defined fibers with Pol III tRNA transcription-associated footprints as those with a footprint between 30 and 140 bp starting 45-55 bp upstream of the TSS, or with a < 90 bp footprint at the B-box or A-box alone (a rare occurrence).

### Defining 5S rRNA Pol III transcription-associated footprints

5S rRNA genes showed a similar pattern of footprints to tRNA genes, but with slightly larger footprints overall. As such, Pol III 5S rRNA transcription associated footprints were defined identically as those found at tRNA genes, except that the maximum size was set to 160 bp (Figure S10).

### Correlation of tRNA expression score to codon frequency

The tRNA expression score was calculated as the fraction of reads containing a Pol III transcription-associated footprint compared to those without. All tRNA genes with coverage within the middle 95% of the overall distribution of coverage across the genome were also identified. Subsequently, all families of tRNA genes sharing a given codon sequence where all known members had an acceptable level of coverage were identified. The Pearson correlation of the overall count of copies of tRNA genes in each family with their codon frequency in the Drosophila genome was then compared to the correlation of codon frequency with the sum of expression scores for each family (Figure S9d).

### Global coordination analysis

The following describes the methodology used for the PPP/PIC coding promoter coordination analysis -- however this approach holds true for all global coordination analyses, substituting the feature (PPP/PIC, promoter accessibility, Pol III transcription associated footprints) and location (Pol II promoter, enhancer, tRNA, 5S rRNA). First, all pairs of genes with a distance between their TSSs of less than 10 kb were identified. Pair of genes were then binned by distance in such a manner as to allow for a roughly equal amount of simultaneous PPP and PIC footprints in each bin. All reads overlapping both members of each pair of genes in each bin were then identified, with the frequency of individual, simultaneous, and absent PPP/PIC counted to generate a contingency table. Only reads where both promoters were accessible were included. Fisher’s exact test was then used to calculate an odds ratio and p-value associated with simultaneous PPP or PIC occupancy in each set of reads binned by distance.

### Per-locus coordination analysis

Coordination on a per-locus level was again quantified identically across features and locations. First, all pairs of genes with a distance between their TSSs of less than 10 kb were identified. All reads overlapping both members of each pair of genes were then identified, with only reads where both promoters were accessible included. The frequency of individual, simultaneous, and absent PPP/PIC footprints for each pair was then counted to generate a contingency table. Fisher’s exact test was then used to identify an associated p-value and odds ratio at each individual locus. Pairs of genes were then assigned as significantly coordinated if the odds ratio was greater than 1 and the p-value was less than 0.05. Gene pairs were then binned based on distance and the fraction of pairs with a significant odds ratio was calculated for each bin.

## Acknowledgements

We thank James Warner, Ines Patop and additional members of the Churchman lab for helpful discussions, assistance, and critical reading of the manuscript; Sean Eddy for discussions about the HMM; K. Munson, and the University of Washington PacBio Sequencing Services for PacBio sequencing. Funding: This work was supported by the NIH (R01-HG007173 to L.S.C., DP5-OD029630 to A.B.S.), and the Brotman Baty Institute for Precision Medicine. A.B.S. holds a Career Award for Medical Scientists from the Burroughs Wellcome Fund and is a Pew Biomedical Scholar. Author Contributions: Conceptualization, T.W.T., L.S.C., and A.B.S.; Methodology, T.W.T. (lead), R.S.I., A.B.S., and L.S.C.; Investigation, T.W.T. (lead), R.S.I., and J.R.; Formal Analysis, T.W.T. (lead), D.D.; Writing - Original Draft, T.W.T..; Writing - Review & Editing, T.W.T., R.S.I., A.B.S, and L.S.C.; Funding Acquisition, A.B.S. and L.S.C.; Supervision, A.B.S. and L.S.C. Competing interests: The authors declare no competing interests. Data and materials availability: Raw and processed sequencing data will be available from the Gene Expression Omnibus (GEO) upon publication. Code for analysis of PacBio sequencing data will be available at GitHub upon publication: www.github.com/church-manlab.

## References

1. Core, L. & Adelman, K. Promoter-proximal pausing of RNA polymerase II: a nexus of gene regulation. Genes Dev. 33, 960–982 (2019).

2. Mayer, A., Landry, H. M. & Churchman, L. S. Pause & go: from the discovery of RNA polymerase pausing to its functional implications. Curr. Opin. Cell Biol. 46, 72–80 (2017).

3. Petesch, S. J. & Lis, J. T. Overcoming the nucleosome barrier during transcript elongation. Trends Genet. 28, 285–294 (2012).

4. Farnung, L., Ochmann, M., Garg, G., Vos, S. M. & Cramer, P. Structure of a backtracked hexasomal intermediate of nucleosome transcription. Mol. Cell 82, 3126–3134.e7 (2022).

5. Wang, H., Schilbach, S., Ninov, M., Urlaub, H. & Cramer, P. Structures of transcription preinitiation complex engaged with the +1 nucleosome. Nat. Struct. Mol. Biol. 30, 226–232 (2023).

6. Chen, Z. et al. High-resolution and high-accuracy topographic and transcriptional maps of the nucleosome barrier. Elife 8, (2019).

7. Ramachandran, S., Ahmad, K. & Henikoff, S. Transcription and Remodeling Produce Asymmetrically Unwrapped Nucleosomal Intermediates. Mol. Cell 68, 1038–1053.e4 (2017).

8. Gilchrist, D. A. et al. Pausing of RNA polymerase II disrupts DNA-specified nucleosome organization to enable precise gene regulation. Cell 143, 540–551 (2010).

9. Yang, L. & Yu, J. A comparative analysis of divergently-paired genes (DPGs) among Drosophila and vertebrate genomes. BMC Evol. Biol. 9, 55 (2009).

10. Herr, D. R. & Harris, G. L. Close head-to-head juxtaposition of genes favors their coordinate regulation in Drosophila melanogaster. FEBS Lett. 572, 147–153 (2004).

11. Small, S. & Arnosti, D. N. Transcriptional enhancers in Drosophila. Genetics 216, 1–26 (2020).

12. Sutherland, H. & Bickmore, W. A. Transcription factories: gene expression in unions? Nat. Rev. Genet. 10, 457–466 (2009).

13. Kimura, H. & Sato, Y. Imaging transcription elongation dynamics by new technologies unveils the organization of initiation and elongation in transcription factories. Curr. Opin. Cell Biol. 74, 71–79 (2022).

14. Patange, S., Ball, D. A., Karpova, T. S. & Larson, D. R. Towards a “Spot On” Understanding of Transcription in the Nucleus. J. Mol. Biol. 433, 167016 (2021).

15. Cisse, I. I. et al. Real-time dynamics of RNA polymerase II clustering in live human cells. Science 341, 664–667 (2013).

16. Cho, W.-K. et al. Mediator and RNA polymerase II clusters associate in transcription-dependent condensates. Science 361, 412–415 (2018).

17. Gressel, S., Schwalb, B. & Cramer, P. The pause-initiation limit restricts transcription activation in human cells. Nat. Commun. 10, 3603 (2019).

18. Farnung, L., Vos, S. M. & Cramer, P. Structure of transcribing RNA polymerase II-nucleosome complex. Nat. Commun. 9, 5432 (2018).

19. Hodges, C., Bintu, L., Lubkowska, L., Kashlev, M. & Bustamante, C. Nucleosomal fluctuations govern the transcription dynamics of RNA polymerase II. Science 325, 626–628 (2009).

20. He, Y., Fang, J., Taatjes, D. J. & Nogales, E. Structural visualization of key steps in human transcription initiation. Nature 495, 481–486 (2013).

21. Krebs, A. R. et al. Genome-wide single-molecule footprinting reveals high RNA polymerase II turnover at paused promoters. Mol. Cell 67, 411–422.e4 (2017).

22. Stergachis, A. B., Debo, B. M., Haugen, E., Churchman, L. S. & Stamatoyannopoulos, J. A. Single-molecule regulatory architectures captured by chromatin fiber sequencing. Science 368, 1449–1454 (2020).

23. Abdulhay, N. J. et al. Massively multiplex single-molecule oligonucleosome footprinting. Elife 9, (2020).

24. Lee, I. et al. Simultaneous profiling of chromatin accessibility and methylation on human cell lines with nanopore sequencing. Nat. Methods 17, 1191–1199 (2020).

25. Shipony, Z. et al. Long-range single-molecule mapping of chromatin accessibility in eukaryotes. Nat. Methods 17, 319–327 (2020).

26. Vos, S. M., Farnung, L., Urlaub, H. & Cramer, P. Structure of paused transcription complex Pol II-DSIF-NELF. Nature 560, 601–606 (2018).

27. Rengachari, S., Schilbach, S., Aibara, S., Dienemann, C. & Cramer, P. Structure of the human Mediator-RNA polymerase II pre-initiation complex. Nature 594, 129–133 (2021).

28. Schilbach, S., Aibara, S., Dienemann, C., Grabbe, F. & Cramer, P. Structure of RNA polymerase II preinitiation complex at 2.9 Å defines initial DNA opening. Cell 184, 4064–4072.e28 (2021).

29. Kwak, H., Fuda, N. J., Core, L. J. & Lis, J. T. Precise maps of RNA polymerase reveal how promoters direct initiation and pausing. Science 339, 950–953 (2013).

30. Hitz, B. C., et al. The ENCODE Uniform Analysis Pipelines. bioRxiv (2023) doi:10.1101/2023.04.04.535623.

31. Luo, Y. et al. New developments on the Encyclopedia of DNA Elements (ENCODE) data portal. Nucleic Acids Res. 48, D882–D889 (2020).

32. ENCODE Project Consortium. An integrated encyclopedia of DNA elements in the human genome. Nature 489, 57–74 (2012).

33. Titov, D. V. et al. XPB, a subunit of TFIIH, is a target of the natural product triptolide. Nat. Chem. Biol. 7, 182–188 (2011).

34. Chereji, R. V., Bryson, T. D. & Henikoff, S. Quantitative MNase-seq accurately maps nucleosome occupancy levels. Genome Biol. 20, 198 (2019).

35. Vos, S. M. et al. Structure of activated transcription complex Pol II-DSIF-PAF-SPT6. Nature 560, 607–612 (2018).

36. Vos, S. M., Farnung, L., Linden, A., Urlaub, H. & Cramer, P. Structure of complete Pol II-DSIF-PAF-SPT6 transcription complex reveals RTF1 allosteric activation. Nat. Struct. Mol. Biol. 27, 668–677 (2020).

37. Shao, W. & Zeitlinger, J. Paused RNA polymerase II inhibits new transcriptional initiation. Nat. Genet. 49, 1045–1051 (2017).

38. Gressel, S. et al. CDK9-dependent RNA polymerase II pausing controls transcription initiation. Elife 6, (2017).

39. Lawrence, J. G. Shared strategies in gene organization among prokaryotes and eukaryotes. Cell 110, 407–413 (2002).

40. Henriques, T. et al. Widespread transcriptional pausing and elongation control at enhancers. Genes Dev. 32, 26–41 (2018).

41. Sartorelli, V. & Lauberth, S. M. Enhancer RNAs are an important regulatory layer of the epigenome. Nat. Struct. Mol. Biol. 27, 521–528 (2020).

42. Mikhaylichenko, O. et al. The degree of enhancer or promoter activity is reflected by the levels and directionality of eRNA transcription. Genes Dev. 32, 42–57 (2018).

43. Arnold, C. D. et al. Genome-wide quantitative enhancer activity maps identified by STARR-seq. Science 339, 1074–1077 (2013).

44. Zabidi, M. A. et al. Enhancer-core-promoter specificity separates developmental and housekeeping gene regulation. Nature 518, 556–559 (2015).

45. Batut, P. J. et al. Genome organization controls transcriptional dynamics during development. Science 375, 566–570 (2022).

46. Wang, Q., Sun, Q., Czajkowsky, D. M. & Shao, Z. Sub-kb Hi-C in D. melanogaster reveals conserved characteristics of TADs between insect and mammalian cells. Nat. Commun. 9, 188 (2018).

47. Beagan, J. A. & Phillips-Cremins, J. E. On the existence and functionality of topologically associating domains. Nat. Genet. 52, 8–16 (2020).

48. Schwartz, Y. B. & Cavalli, G. Three-Dimensional Genome Organization and Function in Drosophila. Genetics 205, 5–24 (2017).

49. Ulianov, S. V. et al. Active chromatin and transcription play a key role in chromosome partitioning into topologically associating domains. Genome Res. 26, 70–84 (2016).

50. Ramírez, F. et al. High-Affinity Sites Form an Interaction Network to Facilitate Spreading of the MSL Complex across the X Chromosome in Drosophila. Mol. Cell 60, 146–162 (2015).

51. Yang, J., Ramos, E. & Corces, V. G. The BEAF-32 insulator coordinates genome organization and function during the evolution of Drosophila species. Genome Res. 22, 2199–2207 (2012).

52. Ong, C.-T., Van Bortle, K., Ramos, E. & Corces, V. G. Poly(ADP-ribosyl)ation regulates insulator function and intrachromosomal interactions in Drosophila. Cell 155, 148–159 (2013).

53. Kaushal, A. et al. CTCF loss has limited effects on global genome architecture in Drosophila despite critical regulatory functions. Nat. Commun. 12, 1011 (2021).

54. Turowski, T. W. & Tollervey, D. Transcription by RNA polymerase III: insights into mechanism and regulation. Biochem. Soc. Trans. 44, 1367–1375 (2016).

55. Graczyk, D., Cieśla, M. & Boguta, M. Regulation of tRNA synthesis by the general transcription factors of RNA polymerase III - TFIIIB and TFIIIC, and by the MAF1 protein. Biochim. Biophys. Acta Gene Regul. Mech. 1861, 320–329 (2018).

56. Male, G. et al. Architecture of TFIIIC and its role in RNA polymerase III pre-initiation complex assembly. Nat. Commun. 6, 7387 (2015).

57. Geslain, R. & Pan, T. Functional Analysis of Human tRNA Isodecoders. J. Mol. Biol. 396, 821–831 (2010).

58. Marygold, S. J., Chan, P. P. & Lowe, T. M. Systematic identification of tRNA genes in Drosophila melanogaster. MicroPubl Biol 2022, (2022).

59. Behrens, A., Rodschinka, G. & Nedialkova, D. D. High-resolution quantitative profiling of tRNA abundance and modification status in eukaryotes by mim-tRNAseq. Mol. Cell 81, 1802–1815.e7 (2021).

60. Lucas, M. C. et al. Quantitative analysis of tRNA abundance and modifications by nanopore RNA sequencing. Nat. Biotechnol. (2023) doi:10.1038/s41587-023-01743-6.

61. Nakamura, Y., Gojobori, T. & Ikemura, T. Codon usage tabulated from the international DNA sequence databases. Nucleic Acids Res. 26, 334 (1998).

62. Procunier, J. D. & Tartof, K. D. Genetic analysis of the 5S RNA genes in Drosophila melanogaster. Genetics 81, 515–523 (1975).

63. Hull, M. W., Erickson, J., Johnston, M. & Engelke, D. R. tRNA genes as transcriptional repressor elements. Mol. Cell. Biol. 14, 1266–1277 (1994).

64. Raab, J. R. et al. Human tRNA genes function as chromatin insulators. EMBO J. 31, 330–350 (2012).

65. Gaertner, B. & Zeitlinger, J. RNA polymerase II pausing during development. Development 141, 1179–1183 (2014).

